# Chronopharmacological targeting of mitochondrial dihydroorotate dehydrogenase prevents diet-induced obesity in male mice

**DOI:** 10.64898/2026.04.18.719366

**Authors:** Florian Atger, Manon Durand, Mikaël Croyal, Albane Chavanne, Enora Le Questel, Yolène Foucher, Charlène Besnard, Ivan Nemazanyy, Soazig Le Lay, Xavier Prieur, Claire Pecqueur, Daniel Mauvoisin, David Jacobi

## Abstract

Daily hepatic mitochondrial rhythms are strengthened by time-restricted feeding in diet-induced obesity in male mice, as shown by metabolomics, including an early enhancement of oscillations within the *de novo* pyrimidine pathway. We tested whether timed inhibition of dihydroorotate dehydrogenase (DHODH, which links pyrimidine synthesis to respiratory-chain flux), can reproduce selected TRF-associated mitochondrial and metabolic effects. Administering the short-half-life inhibitor BAY2402234 at dawn transiently decreased DHODH activity, restored daily mitochondrial oxidative dynamics (ubiquinone/ubiquinol ratio), and amplified rhythms of respiratory exchange ratio and mitochondrial dynamics-related markers upon high-fat diet. Under ZT0 dosing, mice showed reduced weight gain, reduced adiposity and hepatic triglycerides, and improved glucose tolerance without changes in food intake; while ZT12 dosing was ineffective. Hepatic *Dhodh* knockdown did not reproduce the anti-obesity phenotype, and uridine supplementation blunted BAY2402234 benefits, implicating *de novo* pyrimidine flux. Our findings reveal rhythm-aware DHODH inhibition as a chronopharmacological preclinical candidate approach against overnutrition.

---

Obesity affects an estimated 890 million adults worldwide^1^. Modern lifestyle–driven disruption of daily biological rhythms that coordinate metabolic homeostasis is closely linked to obesity^2,3^. Timing of feeding is a critical component of metabolic homeostasis^4–6^, yet high-fat diet (HFD) in mice disrupts feeding rhythms, circadian behavior, and circadian clock-controlled gene expression^7^. Time-restricted feeding (TRF), confining food intake to the active phase, mitigates HFD-induced metabolic dysfunction in mice, improving glucose tolerance^8,9^ and reinforcing rhythmic metabolic programs^9,10^ even when the molecular circadian clock is impaired^11^. These observations suggest that rhythmic downstream effectors, rather than the clockwork alone, could be targeted therapeutically.

Circadian medicine proposes that aligning treatment timing with biological rhythms (chronopharmacotherapy) can enhance efficacy and reduce side effects^12–14^. Although several rodent studies have demonstrated successful chronopharmacological strategies for metabolic disease^15–17^, the application to obesity remains limited, and ongoing clinical efforts emphasize long-acting agents rather than time-specific dosing^18,19^. In particular, whether time-of-day dosing can recapitulate the benefits of feeding-time interventions in obesity remains unknown.

Mitochondria integrate circadian and feeding cues to tune oxidative metabolism over the day–night cycle^20^. Prior work has catalogued daily mitochondrial rhythms across nuclear-encoded genes^21^, protein abundance^22^, redox cofactor cycling (e.g., NAD⁺/NADH)^23–25^, and organelle dynamics^26,27^. Pharmacologically modulating mitochondrial oxidative function can improve metabolic endpoints in diet-induced obesity (DIO)^28–30^. Yet, to our knowledge, no study has tested whether explicitly timing a mitochondrial-targeted intervention can leverage these rhythms to counteract overnutrition. Previous studies link pyrimidine metabolism to mitochondrial dynamics and metabolic homeostasis^31,32^.

Using TRF as a screening strategy, we characterized circadian mitochondrial metabolomic rhythms and identified *de novo* pyrimidine pathways as a candidate therapeutic target in DIO. Beyond supplying nucleotides for replication and RNA/DNA turnover, this pathway is physically and functionally wired into mitochondrial respiration through dihydroorotate dehydrogenase (DHODH), the enzyme that catalyzes the oxidation of dihydroorotate (DHO) to orotate. DHODH is embedded in the inner mitochondrial membrane, where it donates electrons to the ubiquinone (coenzyme Q; CoQ) pool^33,34^. This positioning gives DHODH the potential to influence the CoQ redox state, respiratory control, and the balance between glucose and lipid oxidation, processes that themselves oscillate daily and are reinforced by rhythmic feeding/fasting schedules^31,35–37^. Several features make this coupling attractive for chronopharmacology. First, intermediates flanking DHODH, and the expression of enzymes in the *de novo* pyrimidine pathway, show time-of-day variation and respond to feeding rhythms, indicating that pathway flux is gated by daily physiology^35–37^. Second, because DHODH reduces CoQ, transient changes in its activity could recalibrate the redox poise of the respiratory chain and thereby modulate oxidative capacity without chronic suppression of respiration^33^. Third, nucleotide demand, mitochondrial biogenesis, and macromolecular repair are rhythmic and respond to feeding, creating temporal windows in which brief DHODH engagement might exert outsized effects35–37.

Here, we show that brief inhibition of DHODH during the light-phase realigns diurnal oxidative metabolism and enhances metabolic flexibility in HFD-fed male mice. ZT0 dosing limits weight gain and adiposity and reduces hepatic triglyceride content, whereas identical ZT12 dosing is ineffective. These findings identify time of administration as a key determinant of efficacy and support DHODH as a rhythm-sensitive metabolic target relevant to obesity.

## RESULTS

### Mitochondrial metabolic rhythms are promoted by TRF

To define the temporal relationship between mitochondrial diurnal rhythms and the metabolic benefits of TRF in HFD-fed mice, we compared hepatic and mitochondrial metabolic oscillations after short-term (4 days) and long-term (12 weeks) TRF (Fig. 1a). Short-term TRF did not alter body weight, total energy intake, glucose tolerance, fasting glycaemia, insulin sensitivity, or serum insulin (Extended Data Fig. 1a–f). In contrast, long-term TRF limited HFD-induced weight gain (Extended Data Fig. 1g) without changing cumulative energy intake (Extended Data Fig. 1h), improved glucose tolerance despite unchanged fasting glucose (Extended Data Fig. 1i-j), improved insulin sensitivity (Extended Data Fig. 1k), and lowered serum insulin (Extended Data Fig. 1l), consistent with prior reports^9^. Short-term TRF shifted the phase of core-clock transcripts (*Arntl*, *Nr1d1*, *Per2*) and of the clock-controlled gene *Nampt*, with similar phase shifts after long-term TRF (Extended Data Fig. 1m,n). Consequently, we collected whole-liver lysates and isolated hepatic mitochondria every 4 h across 24 h to profile metabolic oscillations after short- and long-term TRF. We first analyzed metabolite levels irrespective of sampling time using rhythm-adjusted mean (*i.e*. midline estimating statistic of rhythm: mesor) comparison. Short-term TRF produced no detectable remodeling of the liver or mitochondrial metabolomes (Extended Data Fig. 1o). By contrast, long-term TRF altered numerous metabolites (Extended Data Fig. 1p; Supplementary Data 1). With the exception of NAD^+^, which changed in both tissue fractions, significant metabolites were largely compartment-specific, underscoring the value of subcellular profiling to reveal TRF-regulated pathways. We confirmed mitochondrial fraction enrichment, shown by depletion of cytosolic β-actin and nuclear histone H4 in the mitochondrial fraction relative to whole-liver lysates (Extended Data Fig. 1q). In contrast to mesor levels, rhythmicity analyses revealed a distinct pattern: both short- and long-term TRF increased the number of rhythmic metabolites in whole liver and in mitochondria (Fig. 1b). The effect was most pronounced in mitochondria after short-term TRF and remained across a broad range of False Discovery Rate (FDR) cutoffs. At a stringent threshold (FDR = 0.05), TRF increased rhythmic metabolites in mitochondria by >4-fold (short-term) and >2-fold (long-term) relative to *ad libitum*, exceeding the corresponding changes in whole liver (Fig. 1c). To probe how TRF reshapes metabolic oscillations in mitochondria versus whole-liver extracts, we compared rhythmicity patterns across conditions (Fig. 1d,g; Supplementary Data 1). Whole-tissue extracts contained more rhythmic metabolites than mitochondrial fractions (Fig. 1b,e,h), indicating that overall rhythmicity is preserved at the tissue level. In liver, roughly half of rhythmic metabolites were shared between *ad libitum* and TRF (48/98 for short-term; 45/90 for long-term). In mitochondria, rhythmicity was largely TRF-specific (59/71 for short-term; 30/42 for long-term). Peak phase plots revealed an approximately antiphase relationship between compartments, with liver metabolites peaking predominantly in the dark phase and mitochondrial metabolites in the light phase (Fig. 1e,h). Notably, mitochondrial rhythms remained sharply concentrated around ZT6–ZT8 under both short- and long-term TRF, whereas liver peak phases were more dispersed. Class-level phase distributions further highlighted stronger TRF effects in mitochondria than in liver, most prominently an enhancement of rhythmicity in mitochondrial nucleotides across the cycle (Fig. 1f,i). In sum, TRF induces a larger gain in rhythmic metabolites in liver mitochondria than in whole liver, and the distinct peak phase structure indicates that mitochondrial and whole-liver metabolomes display distinct temporal pattern under TRF, without allowing us to conclude that mitochondria respond earlier than the liver as a whole.

**Figure 1:**
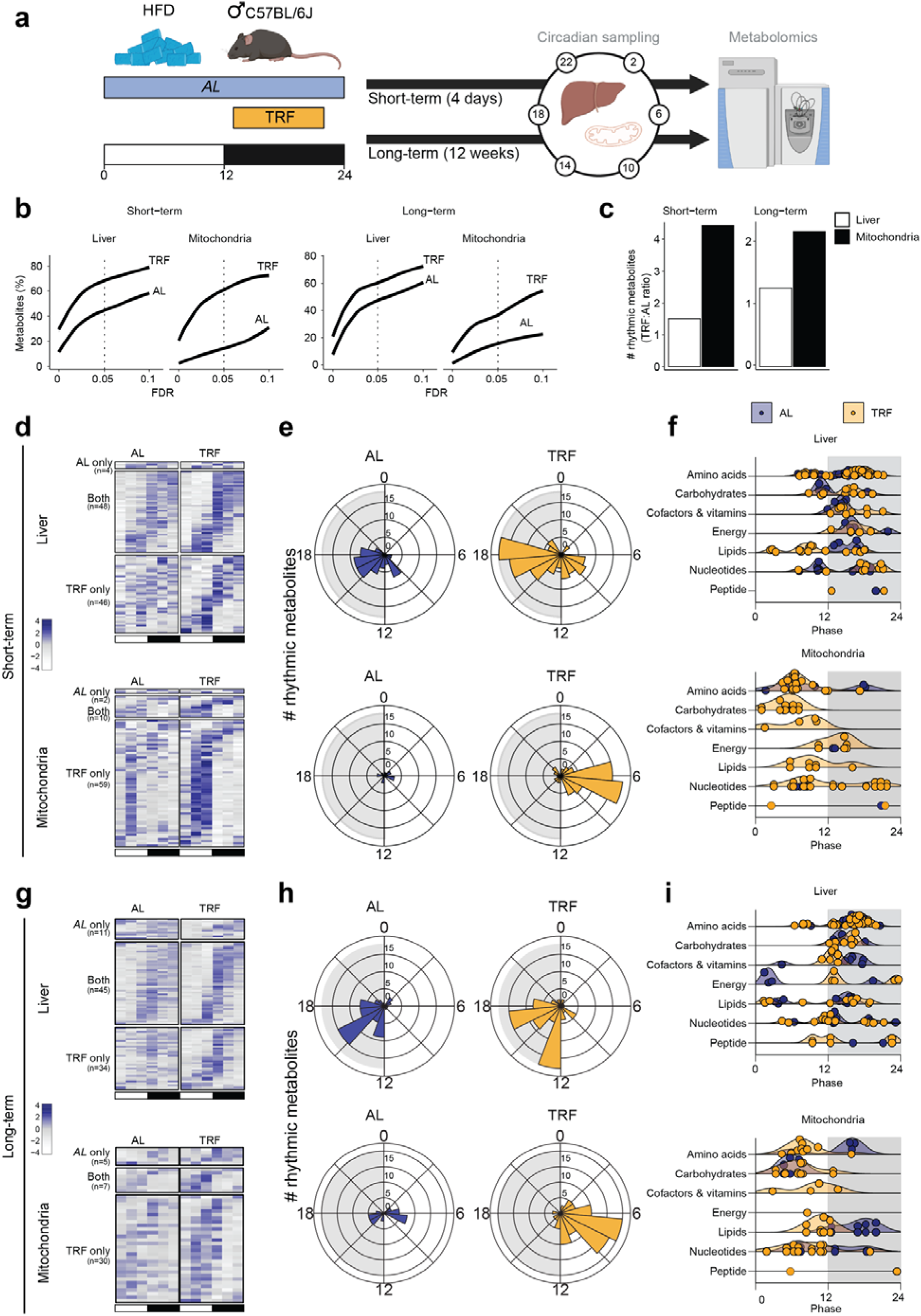
TRF strengthens rhythmicity in the liver mitochondrial metabolome, with persistent class- and phase-specific changes. **a,** Experimental design. Male C57BL/6J mice on HFD were fed *ad libitum* or under time-restricted feeding (TRF) for a short term (4 days) or long term (12 weeks). Liver and isolated hepatic mitochondria were collected every 4 h across 24 h (Zeitgeber time, ZT), and metabolites were analyzed by LC-MS (n = 3-4 mice per ZT per condition). Rhythmic metabolites were identified with CircaCompare^38^. **b,** Cumulative fraction of metabolites called rhythmic in liver or mitochondria as a function of FDR threshold (0–0.1); dotted line marks FDR = 0.05. **c,** TRF/*ad libitum* ratio of the number of rhythmic metabolites at FDR < 0.05 for short- and long-term TRF in liver (white) and mitochondria (black). **d,g,** Heat maps of rhythmic metabolites (FDR < 0.05) in whole-liver extracts (top) and mitochondria fractions (bottom) for short-term **(d)** and long-term **(g)** experiments. Metabolites are grouped as rhythmic in *ad libitum* only, in both conditions, or in TRF only. Black/white bars indicate dark/light phases. **e,h,** Rose plots showing phase distribution of rhythmic metabolites in liver (top) and mitochondria (bottom) for short-term **(e)** and long-term **(h)** experiments. **f,i,** Phase distribution by metabolite class for short-term **(f)** and long-term **(i)** experiments. Points denote individual metabolites; shaded densities summarize class-level phase preference (shown only when ≥4 metabolites in a class are rhythmic). See also Extended Data Fig. 1 and Supplementary Data 1.

### TRF enhances daily rhythmicity in mitochondrial oxidative functions

To test whether TRF promotes rhythmicity of mitochondrial pathways, we analyzed transcript dynamics alongside metabolites. In line with the metabolome, both short- and long-term TRF increased the number of rhythmic transcripts (Extended Data Fig. 2a) and, in the long-term condition, showed high overlap with a comparable dataset (Deota *et al.* 2023; *ad libitum* versus TRF, 10-week HFD; Extended Data Fig. 2b^39^). Gene Ontology analysis for cellular components revealed enrichment of mitochondria-related terms among TRF-induced rhythmic transcripts (Extended Data Fig. 2c). Among mitochondria-annotated rhythmic transcripts, peak phases shifted predominantly to the dark period and amplitudes increased under TRF (Extended Data Fig. 2d). Notably, TRF induced rhythmicity in nuclear-encoded genes central to mitochondrial respiration (Extended Data Fig. 2e). An integrative transcriptomic–metabolomic analysis showed that both short- and long-term TRF specifically enhance rhythmicity in pathways linked to mitochondrial oxidative processes and energy metabolism (FDR < 0.05; Supplementary Data 2; Extended Data Fig. 2f,g). These findings point to improved metabolic flexibility under TRF, *i.e*., a coordinated daily shift between carbohydrate and lipid oxidation, likely driven by mitochondrial rhythmicity. TRF strengthened rhythms in glycolytic and pyruvate-pathway metabolites, which support mitochondrial redox (NAD⁺/NADH cycling) and acetyl-CoA supply to the TCA cycle (Extended Data Fig. 2h). It also enhanced rhythmicity of TCA intermediates and of nicotinamide metabolism (Extended Data Fig. 2i), pathways known to regulate mitochondrial function under circadian and feeding/fasting control^23,24^. TRF further affected glutathione (GSH), a key mitochondrial antioxidant: synthesized in the cytosol, GSH is imported into mitochondria where oxidation to glutathione disulfide (GSSG) detoxifies reactive oxygen species^40^ (Extended Data Fig. 2h). TRF induced rhythmic patterns in the GSH:GSSG and succinate:fumarate ratios, while concurrently reducing the lactate to pyruvate ratio (Extended Data Fig. 2h,i; Supplementary Data 3), consistent with TRF-dependent remodeling of redox-related metabolic rhythms.

### TRF promotes rhythms in mitochondrial oxidative metabolism and *de novo* pyrimidine biosynthesis pathways

Through untargeted profiling across short-term and long-term TRF, we observed a consistent enhancement in mitochondrial metabolic rhythms. Three of the most promoted rhythms map to the *de novo* pyrimidine biosynthesis: carbamoyl aspartate, DHO, and orotate (Fig. 2a,b). These metabolites peak in the dark phase; short-term TRF amplifies this pattern (Fig. 2c) and it persists with long-term TRF (Fig. 2d; Supplementary Data 1). Although most steps of *de novo* pyrimidine synthesis are cytosolic, the conversion of DHO to orotate is catalyzed by DHODH in the inner mitochondrial membrane, which uses coenzyme Q (CoQ) as the electron acceptor, thereby coupling pyrimidine synthesis to the respiratory chain^41^ (Fig. 2e). To probe this coupling, we examined the oxidized-to-reduced ratio CoQ9/CoQ9H₂ as a proxy of DHODH activity: it exhibited stronger daily variation under TRF, in phase with pyrimidine-pathway rhythms (Fig. 2f; Supplementary Data 2). Notably, this effect was not due to diurnal changes in DHODH protein abundance (Fig. 2g). These findings collectively suggest that TRF accentuates mitochondrial oxidative function and highlight DHODH-linked pyrimidine metabolism as one TRF-sensitive component of this remodeling.

**Figure 2:**
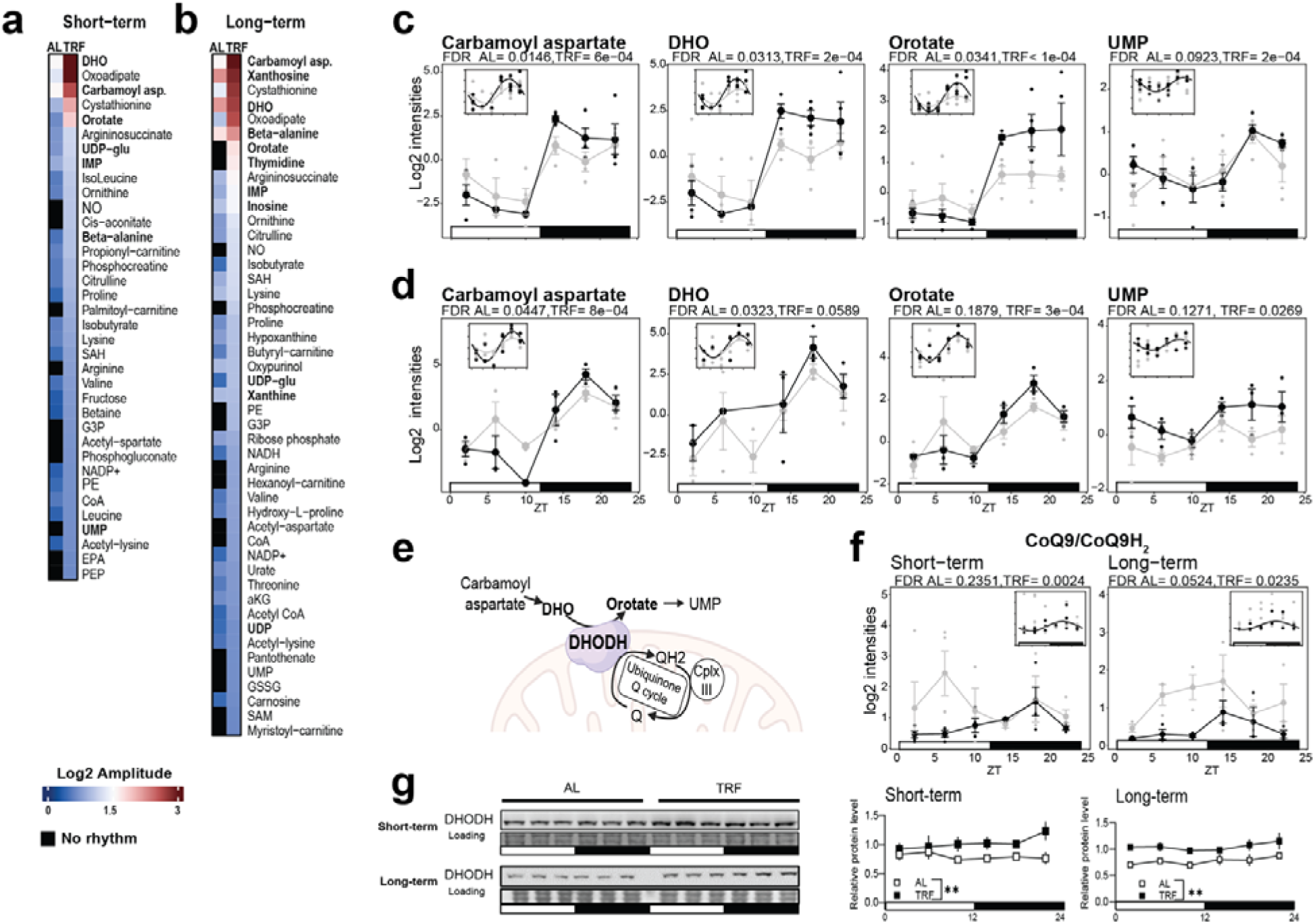
TRF promotes rhythms in mitochondrial oxidative metabolism and *de novo* pyrimidine biosynthesis. **a,b,** Heat maps of rhythmic metabolite amplitude (log2) (FDR < 0.05) in mouse liver for short-term (**a**) and long-term (**b**) experiments (*ad libitum* versus TRF). **c,d,** Time courses (mean ± s.e.m., n = 3–4 mice per ZT per condition) for liver metabolites upstream and downstream of DHODH in short-term (**c**) and long-term (**d**) experiments. CircaCompare FDR for each condition is shown in the panels. **e**, Schematic of the coupling between mitochondria and the *de novo* pyrimidine biosynthesis pathway via DHODH (inner mitochondrial membrane). **f**, Temporal variations in CoQ9/CoQ9H_2_ (ubiquinone:ubiquinol) ratio (mean ± s.e.m., n = 3–4 per ZT per condition). **g,** Immunoblots of liver DHODH across ZT in *ad libitum* and TRF mice for short- and long-term experiments; right, quantification. Naphtol Blue Black staining (LC, loading control) was used for normalization. Experimental design shown in Fig. 1a. Statistical tests: rhythmicity by CircaCompare (panels c–d,f); DHODH levels by two-way ANOVA (* P < 0.05 and ** P < 0.01 where indicated). All experiments were performed in male C57BL/6J mice.

Because TRF modestly elevates hepatic DHODH protein abundance (Fig. 2g), we next tested whether DHODH itself modulates metabolic outcomes using a liver-specific *Dhodh* knockdown (KD), confirmed by immunoblot (Fig. 3a). DHODH KD did not alter body weight upon HFD (Fig. 3b) but improved glucose tolerance (lower glycaemia during glucose tolerance test (GTT) and reduced AUC; Fig. 3c,d), indicating better glucose homeostasis versus controls. To assess effects on pyrimidine biosynthesis, we quantified key intermediates in serum and liver (Fig. 3e). DHO and orotate were unchanged in both compartments. Liver uridine was also unchanged, whereas serum uridine increased with DHODH KD, consistent with a compensatory adjustment in pyrimidine metabolism.

**Figure 3:**
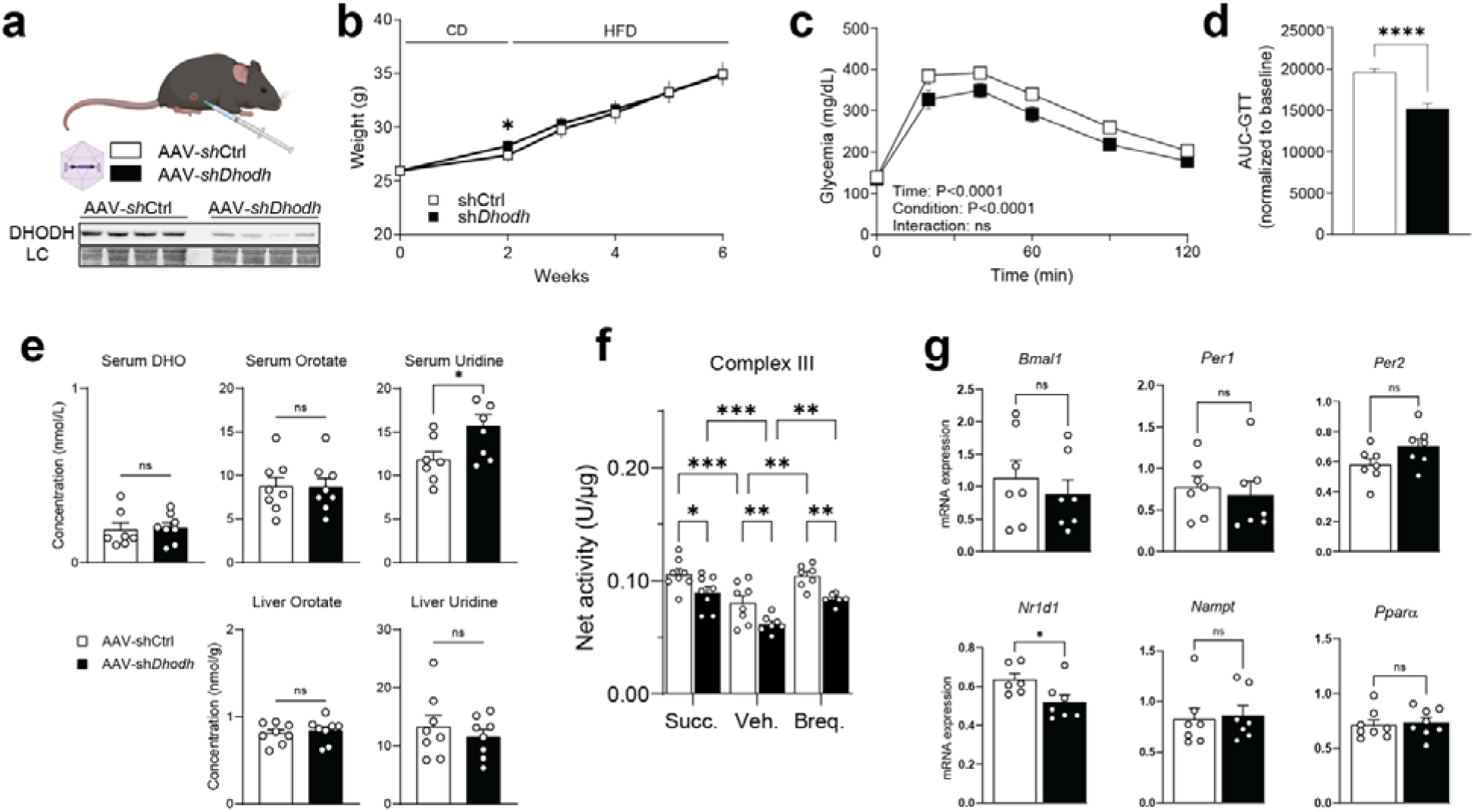
Liver-specific *Dhodh* knockdown improves glucose tolerance and alters mitochondrial/circadian readouts in HFD-fed mice. **a,** Experimental design (top) and immunoblot validation (bottom) of AAV8-mediated liver *Dhodh* knockdown (AAV-sh-*Dhodh*) versus control (AAV-sh-Ctrl). LC, loading control. **b,** Body-weight trajectories under chow (CD) then high-fat diet (HFD); mean ± s.e.m., n = 8 mice per group. **c,** Glucose tolerance test (GTT); two-way ANOVA shows main effects of time and condition (values in panel), interaction ns. **d,** GTT AUC normalized to baseline; n = 8 mice per group. **e,** Serum and liver concentrations of dihydroorotate (DHO), orotate, and uridine; n = 8 mice per group. **f,** Complex III activity in liver homogenates under succinate drive (Succ.), vehicle (Veh.), or brequinar (Breq.) treatment; n = 5 mice per group. **g,** Hepatic mRNA expression of circadian/metabolic genes (*Bmal1*, *Per1*, *Per2*, *Nr1d1*, *Nampt*, *Ppar*α). Statistics: mean ± s.e.m.; ns, not significant; *P < 0.05, **P < 0.01, ***P < 0.001, ****P < 0.0001 (two-way ANOVA with post hoc tests where applicable). All experiments were performed in male C57BL/6J mice.

DHODH KD caused a global reduction in complex III activity (Fig. 3f). Nevertheless, the similar responses to succinate-driven stimulation and to brequinar in control and sh-*Dhodh* mice show that, while diminished, OXPHOS is not abolished under DHODH KD (Fig. 3f). We next profiled hepatic transcripts linked to circadian and metabolic output (*Bmal1*, *Per1*, *Per2*, *Nr1d1*, *Nampt*, *Ppar*α; Fig. 3g) and found that *Nr1d1* was the only transcript significantly reduced by DHODH KD, pointing to a potential crosstalk between DHODH activity and circadian gene regulation.

To test whether DHODH modulates TRF responses, we induced liver-specific *Dhodh* KD, and, two weeks later, fed mice HFD under *ad libitum* or TRF for 4 weeks. *Dhodh* KD did not affect body-weight gain in either regimen, indicating that weight differences were driven by TRF (Extended Data Fig. 3a). TRF improved glucose tolerance, and adding *Dhodh* KD conferred no additional benefit (Extended Data Fig. 3b). Liver orotate levels over time were unchanged by *Dhodh* KD during TRF (Extended Data Fig. 3c). *Dhodh* KD was confirmed longitudinally by qPCR (Extended Data Fig. 3d) but hepatic expression of *Bmal1, Per1, Per2, Nr1d1, Nampt,* and *Ppar*α remained similar to TRF control (Extended Data Fig. 3e). Thus, under HFD, the benefits of TRF on body weight and glucose tolerance appear independent of hepatic *Dhodh* expression.

### Time-of-day DHODH inhibition promotes mitochondrial oxidative metabolism and elongates mitochondria

Because TRF strengthens mitochondrial rhythmicity and DHODH-related metabolites, we hypothesized that transient DHODH inhibition with BAY2402234, a short-half-life DHODH inhibitor^42^, during the light (fasting) phase (ZT0) would synchronize hepatic mitochondrial oxidative metabolism with the fasting-related energy requirements of the liver and peripheral organs during that period.

We first confirmed that BAY2402234 had comparable pharmacokinetics after ZT0 or ZT12 injection (t½ ≈ 4–6 h) across serum, liver, eWAT and brain (Extended Data Fig. 4a,b), supporting timed dosing. We then put HFD-fed mice on a 4-day protocol in which they received vehicle or BAY2402234 at ZT0 or ZT12, in parallel with a short-term TRF schedule (Fig. 4a). Timed DHODH inhibition inverted the temporal profiles of hepatic DHO and orotate when BAY2402234 was given at ZT0 and blunted the rate of change of orotate when given at ZT12 (Fig. 4b, Extended Data Fig. 4c). Circulating DHO and orotate concentrations across time are presented in Extended Data Fig. 4d). ZT0 BAY2402234 also induced time-of-day variations in the CoQ9/CoQ9H_2_ ratio (Fig. 4c), and respiratory control ratio measurements showed improved mitochondrial efficiency after ZT0, but not ZT12, treatment (Fig. 4d). In line with this, ZT0 BAY2402234 increased the ATP:ADP ratio in a time-dependent manner (Fig. 4e), supporting the view that chronopharmacological DHODH inhibition promotes rhythmic mitochondrial oxidation in HFD.

**Figure 4:**
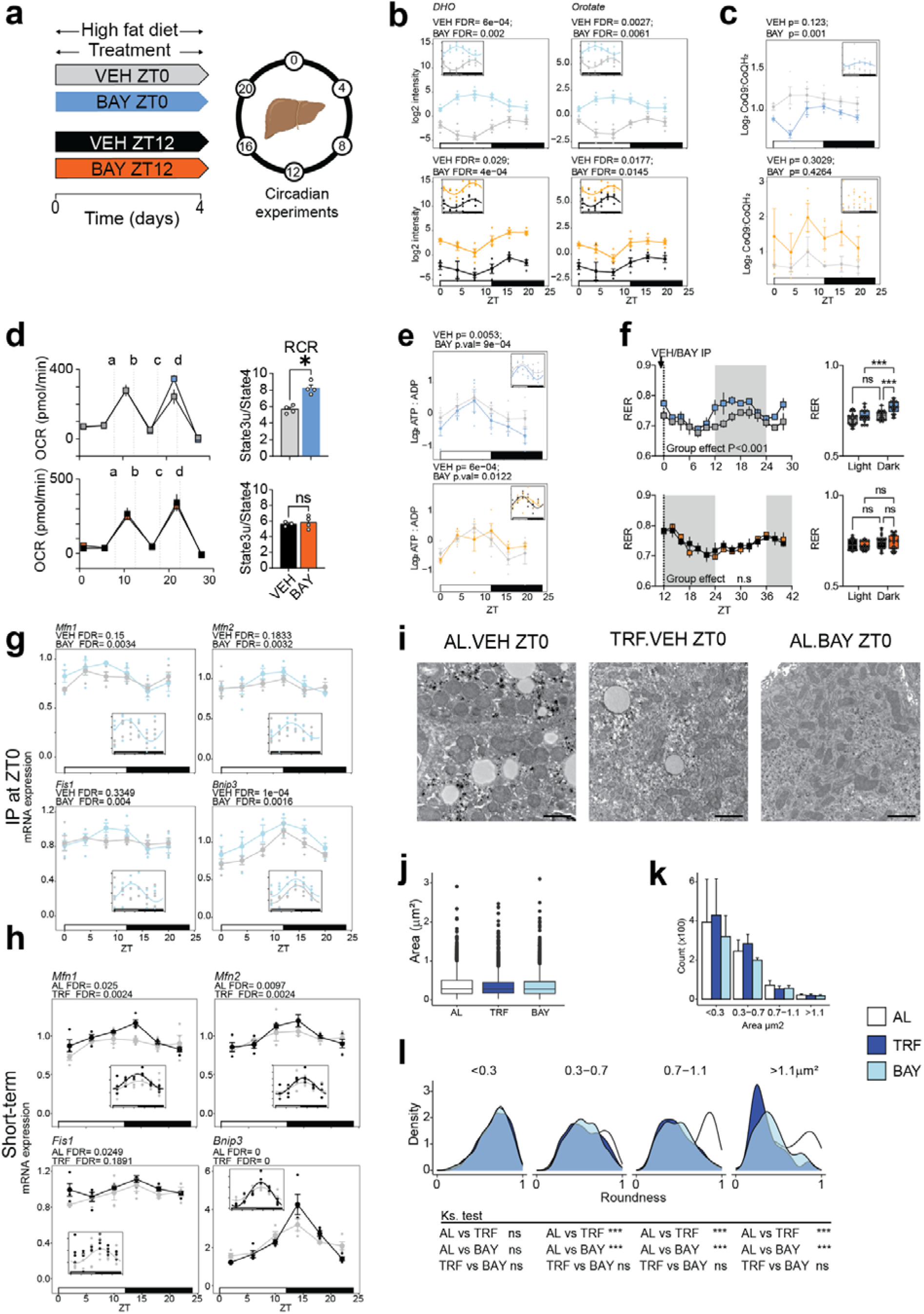
Timed inhibition of DHODH promotes daily mitochondrial oxidative metabolism and elongates mitochondria. **a,** Experimental design. Mice were fed a high-fat diet (HFD) and received daily IP injections of vehicle (VEH) or BAY2402234 at ZT0 or ZT12 for 4 days; livers were collected every 4 h after the last injection. **b,** Hepatic DHO (left) and orotate (right) levels across the 24-h cycle in mice treated at ZT0 (blue) or ZT12 (orange) (n = 3–4 per ZT per condition). **c,** Temporal variations in the CoQ9/CoQ9H₂ ratio in liver (n = 3–4 per ZT per condition). **d,** Oxygen consumption rate measured in isolated mitochondria from mouse livers (n=4 mice/condition). Liver mitochondria were isolated 4 hours following a single injection of BAY2402234 at ZT0 or at ZT12 upon chow diet. Letters a–d indicate sequential additions of ADP, oligomycin, FCCP and antimycin A. Right, respiratory control ratio (RCR) quantification (n = 4 per condition). **e,** Hepatic ATP:ADP ratio across the 24-h cycle after vehicle or BAY2402234 dosing at ZT0 or ZT12 (n = 3–4 per ZT per condition). **f,** Respiratory exchange ratio (RER) in metabolic cages after a single BAY2402234 injection at ZT0 (top) or ZT12 (bottom) in HFD-fed mice; right, RER during light and dark phases (n = 3 per condition). **g,** Expression (RT–qPCR) of genes involved in mitochondrial dynamics (*Mfn1, Mfn2, Fis1, Bnip3*) in HFD-fed mice injected at ZT0 with VEH or BAY2402234 for 4 days (n = 3–4 per ZT per condition). **h,** Expression of the same mitochondrial-dynamics genes in mice subjected to short-term TRF (n = 3–4 per ZT per condition). **i,** Representative transmission electron microscopy images of liver sections from *ad libitum* VEH ZT0 (AL.VEH ZT0), TRF VEH ZT0 (TRF.VEH ZT0) and *ad libitum* BAY2402234 ZT0 (AL.BAY ZT0). Scale bars, 1 μm. **j,** Quantification of mitochondrial area from TEM images in the three groups (n = 3–4 mice per condition). **k,** Mitochondrial size distribution in the three groups (n = 3–4 mice per condition). **l,** Density plots of mitochondrial roundness stratified by area class (<0.3; 0.3–0.7; 0.7–1.1; >1.1 μm²) for each group, with Kolmogorov–Smirnov tests below (AL versus TRF, AL versus BAY, TRF versus BAY). Mitochondrial parameters were quantified from TEM images covering ∼1,500 μm² of liver per mouse. Data are mean ± s.e.m.; *P < 0.05, **P < 0.01, ***P < 0.001. All experiments were performed in male C57BL/6J mice.

To test whether this translated into whole-body fuel flexibility, we acutely inhibited DHODH at ZT0 or ZT12 in mice on a 3-day HFD and performed indirect calorimetry. At day 4, only ZT0 BAY2402234 increased the nocturnal respiratory exchange ratio (RER), indicating enhanced nighttime carbohydrate use, whereas ZT12 dosing had no effect (Fig. 4f).

Because these effects pointed to a broader, liver-wide reorganization of metabolic rhythms, we profiled hepatic metabolites over 24 h after ZT0 or ZT12 BAY2402234. ZT12 dosing reduced the number of rhythmic metabolites compared to controls, whereas ZT0 dosing shifted phase expression (Extended Data Fig. 4e,f). Most metabolites affected by BAY2402234 mapped to circadian-regulated pathways, including nucleotides and mitochondrial substrates (Supplementary Data 4), and circadian core clock gene expression was unchanged (Extended Data Fig. 4g). The time-dependent effects of BAY2402234 on metabolism likely involve mechanisms that extend beyond the molecular circadian clock. UpSet analysis showed that changes in mean liver levels were largely condition-specific (Extended Data Fig. 4h), while rhythmicity analysis revealed that TRF gained the largest number of rhythms, BAY at ZT12 lost many of them, and BAY at ZT0 gained only a subset (Extended Data Fig. 4i). Thus, ZT0 dosing yielded more gained or retained rhythms than ZT12, underscoring a clear time-of-day dependence.

The hepatic clock regulates mitochondrial dynamics (fusion, fission, mitophagy) to adapt to nutrient availability, and disrupting these processes promotes HFD-induced defects^26,43–45^. Because DHODH inhibition was reported to elongate mitochondria in mouse cells^31^, we asked whether ZT0 BAY2402234 would similarly influence mitochondrial remodeling. ZT0 treatment increased the rhythmic expression of genes controlling mitochondrial shape (Fig. 4g), recapitulating the pattern seen with short-term TRF (Fig. 4h). Electron microscopy at ZT4 (4 h after ZT0 dosing) showed that neither ZT0 BAY2402234 nor short-term TRF altered mitochondrial area or size distribution (Fig. 4i–k), but ZT0 BAY2402234 elongated mitochondria and corrected the bulbous morphology observed in *ad libitum* HFD livers (Fig. 4l), again paralleling TRF.

Taken together, light-phase (ZT0/lights-on) chronopharmacological inhibition of DHODH in HFD-fed mice rapidly alters mitochondrial function, metabolite rhythmicity and mitochondrial morphology in a manner that partially recapitulates short-term TRF, whereas ZT12 dosing is not effective.

### Timing of DHODH inhibition determines transient mitochondrial metabolome remodeling in DIO mice

To test whether the metabolic benefits of time-dependent DHODH inhibition in HFD-fed mice are accompanied by longer-lasting changes in hepatic mitochondrial oxidative metabolism, we used mice maintained on a 12-week HFD. During the last 4 weeks, animals received IP injections of vehicle (VEH) or BAY2402234 every other day, so as to limit residual inhibition across two consecutive light/dark cycles, following a chronopharmacology approach previously described^15^. Liver mitochondria were isolated 4 h and 16 h after the final injection at ZT0 or ZT12 (Fig. 5a).

**Figure 5:**
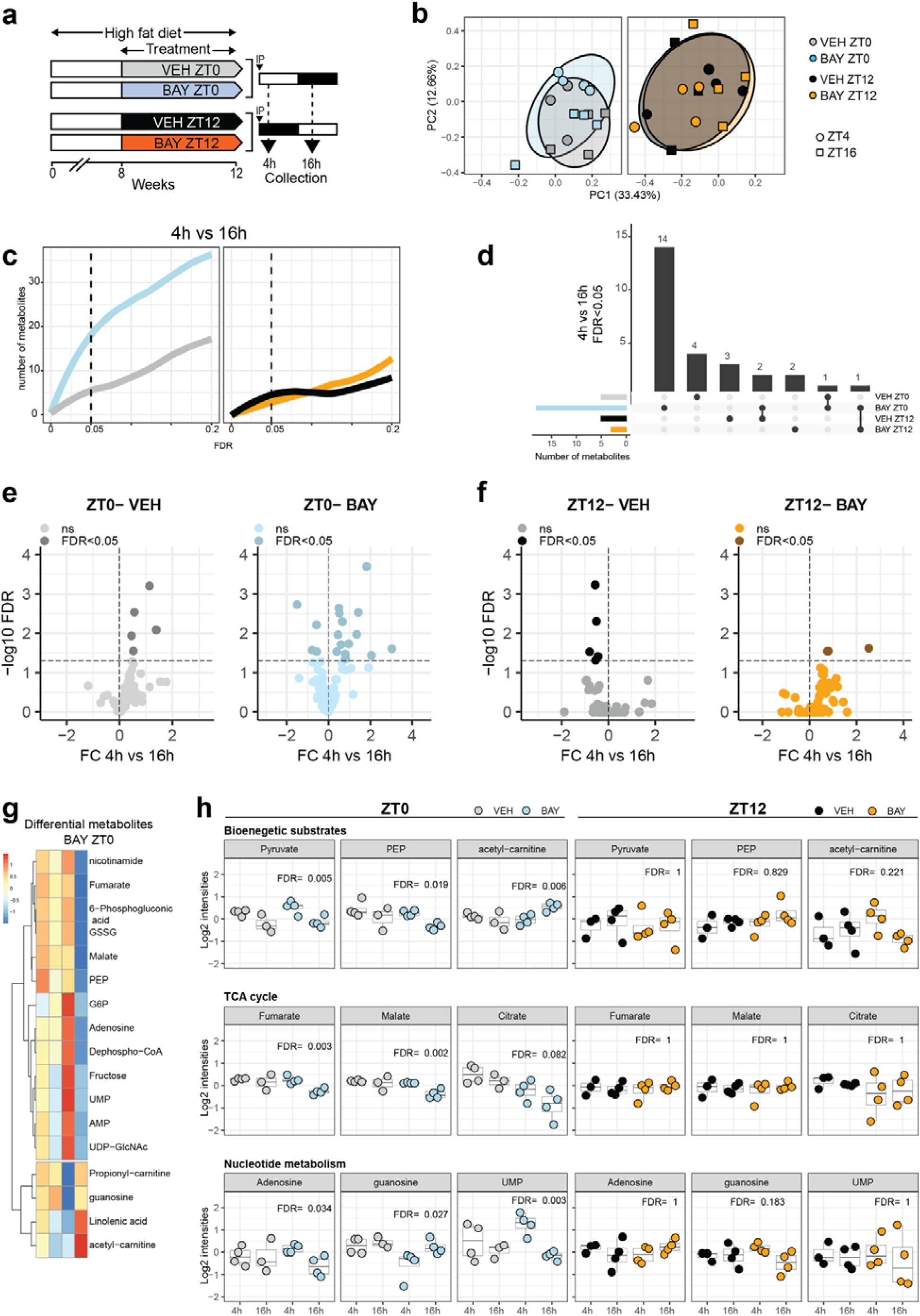
Time-of-day DHODH inhibition differentially remodels the hepatic mitochondrial metabolome in HFD-fed mice. **a,** Experimental design. Mice were fed a high-fat diet (HFD) for 12 weeks. During the last 4 weeks they received IP injections of vehicle (VEH) or BAY2402234 at ZT0 or ZT12 on alternate days. Livers were collected 4 h (ZT4) or 16 h (ZT16) after the final injection, and liver mitochondria were isolated for metabolomics (n = 4 mice per group). **b,** Principal component analysis (PCA) of mitochondrial metabolites. Samples do not separate by treatment (VEH versus BAY) but show an additional separation by sampling time (ZT4 versus ZT16) specifically in the BAY ZT0 group. **c,** Cumulative number of mitochondrial metabolites showing a difference between 4 h and 16 h across groups as a function of FDR. The vertical dashed line indicates FDR = 0.05. Note the greater number of time-regulated metabolites after BAY at ZT0 (blue) than after BAY at ZT12 (orange). **d,** UpSet plot of metabolites that differ between 4 h and 16 h at FDR < 0.05 across the four groups (VEH-ZT0, BAY-ZT0, VEH-ZT12, BAY-ZT12). A unique set of 14 metabolites is specific to BAY-ZT0. **e,** Volcano plots showing fold change (FC, 4 h versus 16 h post IP) versus –log10 FDR for mitochondrial metabolites in ZT0 groups: VEH-ZT0 (left) and BAY-ZT0 (right). BAY at ZT0 increases the number of metabolites with significant time-dependent variation. **f,** Same analysis for ZT12 groups: VEH-ZT12 (left) and BAY-ZT12 (right). BAY at ZT12 has a much weaker effect on time-dependent metabolite variation. **g,** Heatmap of the 14 time-regulated metabolites specific to BAY ZT0, including bioenergetic substrates and TCA-cycle intermediates. **h,** Time-course plots (4 h versus 16 h post IP) for representative metabolites grouped by pathway, bioenergetic substrates (pyruvate, PEP, acetyl-carnitine), TCA cycle (fumarate, malate, citrate), and nucleotide metabolism (adenosine, guanosine, UMP), comparing VEH and BAY injected at ZT0 (left panels, blue) or at ZT12 (right panels, orange). FDR values are indicated for each metabolite. All experiments were performed in male C57BL/6J mice.

The strongest remodeling of the mitochondrial metabolome was observed when BAY2402234 was given at ZT0, particularly at 4 h post-dose. ZT0 treatment elicited rhythmicity in a larger set of mitochondrial metabolites than ZT12 at both 4 h and 16 h (Extended Data Fig. 5a–c). Comparing altered metabolites between conditions and time points showed mostly condition-specific signatures, arguing against a sustained mitochondrial remodeling regardless of dosing time (Extended Data Fig. 5c). Metabolites modified after ZT0 dosing included intermediates of mitochondrial bioenergetics (NAD⁺, NADP⁺, ADP, ATP) and of the TCA cycle (malate, fumarate, citrate, cis-aconitate), each with a distinct temporal profile (Extended Data Fig. 5d,e), whereas the mitochondrial metabolome of mice dosed at ZT12 remained more similar to their controls (Extended Data Fig. 5f,g).

### BAY2402234 treatment at ZT0 triggers temporal variation in the mitochondrial metabolome of DIO mice

We next asked whether BAY2402234 could restore daily variation of the mitochondrial metabolome in DIO mice. Principal component analysis (PCA) showed that, overall, mitochondrial profiles from BAY2402234-treated mice did not segregate from vehicle controls at either dosing time. By contrast, an additional separation according to sampling time (ZT4 versus ZT16) became apparent when BAY2402234 was given at ZT0, but not when it was given at ZT12 (Fig. 5b). Consistent with this, comparison of mitochondrial metabolite levels at 4 h versus 16 h post-dose indicated that only ZT0 injection increased the number of differentially time-regulated metabolites (Fig. 5c). Intersection analysis across all groups identified a distinct signature specific to BAY2402234 at ZT0, comprising 14 unique time-regulated metabolites (Fig. 5d). Accordingly, the ZT4-to-ZT16 fold-change pattern imprinted by BAY2402234 at ZT0 was clearly different from that observed after ZT12 dosing (Fig. 5e,f). The metabolites driving this pattern mapped to canonical rhythmic mitochondrial pathways, including bioenergetic substrates (pyruvate, PEP, acetyl-carnitine), TCA-cycle intermediates (fumarate, malate, citrate) and nucleotide metabolism (adenosine, guanosine, UMP), all showing differential 4 h versus 16 h levels (Fig. 5g,h).

These findings indicate that BAY2402234 engages DHODH at both times of day, but only ZT0 dosing imprints a robust, time-of-day–dependent remodeling of the hepatic mitochondrial metabolome. In mice on long-term HFD, ZT0 administration selectively reinstated temporal differences between ZT4 and ZT16 in mitochondrial bioenergetics and TCA-cycle intermediates, whereas ZT12 dosing left the mitochondrial metabolome largely similar to vehicle. Thus, timed (ZT0) DHODH inhibition enhances the coordination of glycolysis/pyruvate use, TCA flux and lipid-related pathways with the light/dark cycle, consistent with improved metabolic alignment.

### Transient DHODH inhibition with BAY2402234 at ZT0 prevents diet-induced obesity in mice

We next tested whether the light-phase window identified above (injection at ZT0) was sufficient to improve whole-body metabolism in mice on a 12-week HFD (Fig. 6a). BAY2402234 given at ZT0 prevented HFD-induced body-weight gain (Fig. 6b,c) and significantly reduced fat mass (Fig. 6d). Liver weight was unchanged (Fig. 6e), but hepatic triglyceride content was lowered (Fig. 6f). ZT0 treatment also improved insulin sensitivity, as shown by better glucose tolerance (Fig. 6g,h) and lower fasting glycemia (Fig. 6i). These effects occurred without changes in total food intake (Fig. 6j) or in its light/dark distribution (Fig. 6k). None of the benefits seen with BAY2402234 timed at ZT0 were observed when the drug was administered at ZT12 (Extended Data Fig. 6a-k).

**Figure 6:**
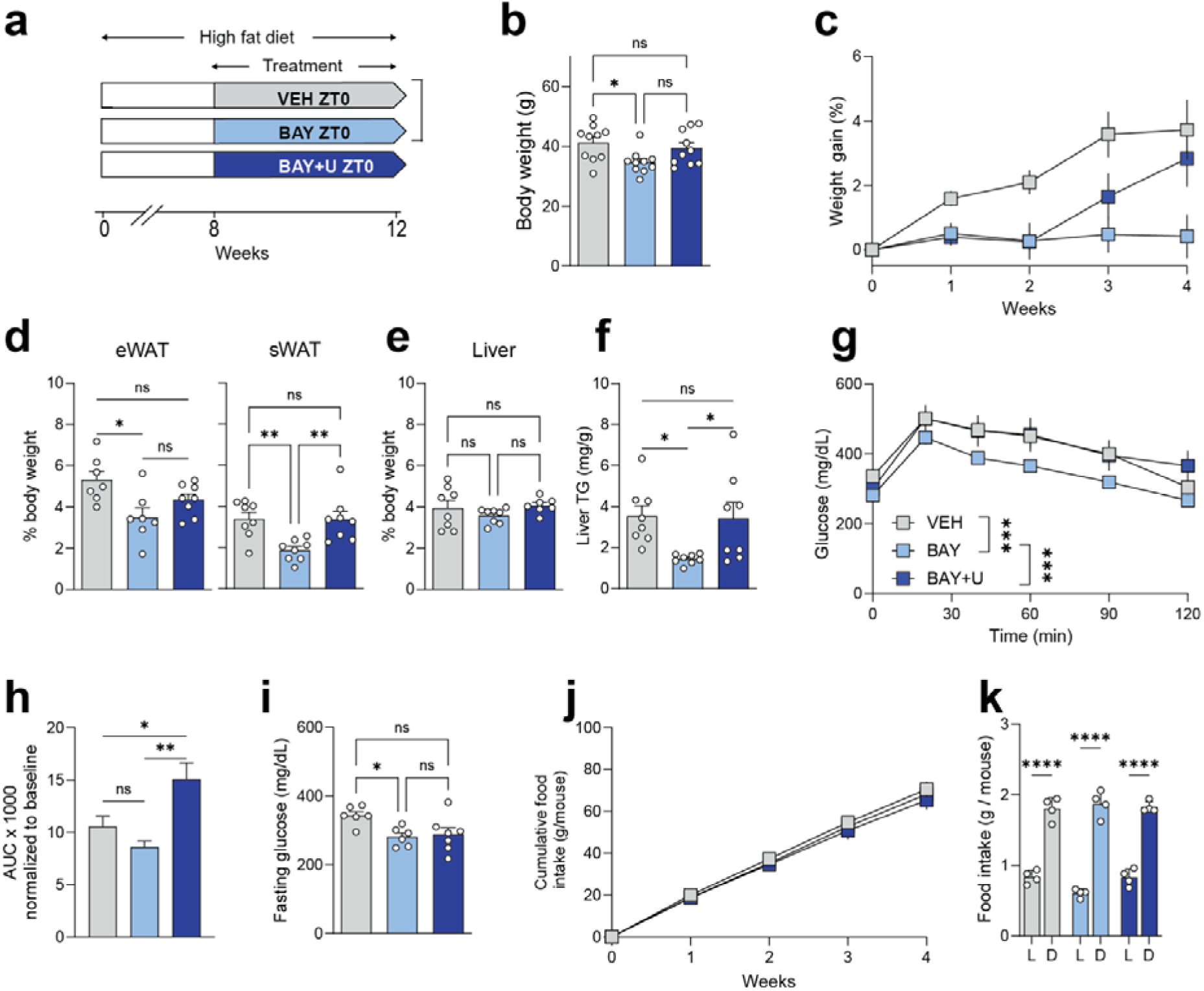
DHODH inhibition at ZT0 prevents diet-induced obesity in male mice without altering food intake. **a,** Experimental design. Mice were fed a HFD for 12 weeks. From week 8 to week 12 they received IP injections every other day at ZT0 of vehicle (VEH), BAY2402234 (BAY, 1.6 mg/kg), or BAY2402234 together with uridine in the drinking water (BAY+U, 5 g/L) (n = 10/group). **b,** Terminal body weight (n=10/group). **c,** Body-weight gain (%) during the 4-week treatment (n = 10/group). **d,** Epididymal (eWAT) and subcutaneous (sWAT) fat mass expressed as % body weight (n = 9–10/group). **e,** Liver weight as % body weight (n = 9–10/group). **f,** Hepatic triglyceride (TG) content (n = 8/group). **g,** Glucose tolerance test (GTT) at week 3, 6 h after the last injection; mice were fasted for 6 h before the test (n = 6/group). **h,** GTT area under the curve (AUC) corresponding to panel g. **i,** Fasting blood glucose (n = 6/group). **j,** Cumulative food intake per mouse over the 4-week treatment (n = 10/group). **k,** Food intake during the light (L) and dark (D) phases (3 independent 24-h measurements per group over the 4 weeks). Data are mean ± s.e.m. ns, not significant; *P < 0.05; **P < 0.01; ***P < 0.001; ****P < 0.0001. One-way ANOVA (or Welch’s t-test, as appropriate) was used for panels b, d–f, h–k; two-way ANOVA with Bonferroni’s post-hoc test was used for panels c and g. See also Extended Data Fig. 6 for ZT12-treated conditions. All experiments were performed in male C57BL/6J mice.

To determine whether these timed effects of BAY2402234 required interference with pyrimidine biosynthesis, we co-administered uridine. Uridine did not alter the early body-weight benefit of ZT0 BAY2402234, but weight gain resumed during the second half of treatment (Fig. 6c), and the reduction in subcutaneous white adipose tissue (sWAT) mass was no longer evident at the end point (Fig. 6d). Uridine also blunted the BAY2402234-induced decrease in liver triglycerides (Fig. 6f). Moreover, in the GTT, uridine abolished the glucose-tolerance benefit conferred by BAY2402234 and brought responses back to, or slightly above, VEH levels (Fig. 6g,h).

These reversals occurred despite similar cumulative and daily food intake across groups (Fig. 6j,k). Because these data suggested that the anti-obesity effect was not due to reduced feeding, we increased the injection frequency from every other day to daily for one week. Food intake and its circadian distribution remained unchanged, irrespective of whether injections were at ZT0 or ZT12 (Extended Data Fig. 7a,b), confirming that BAY2402234 is not anorexigenic in this setting. Anorexigenic signals (hepatic FGF21, circulating GDF15) were unchanged after ZT0 dosing but rose 16 h after ZT12 dosing (Extended Data Fig. 7c–h), indicating that the inactive time window is the one that more strongly induces stress/anorexigenic pathways.

Taken together, these findings show that only light-phase (ZT0) chronopharmacological inhibition of DHODH prevents HFD-induced weight gain and improves glucose–lipid homeostasis without reducing caloric intake, and that part of this protection is sensitive to uridine supplementation, pointing to the pyrimidine biosynthetic pathway as one contributor.

## DISCUSSION

Time-restricted feeding (TRF) in HFD-fed male mice strengthens daily mitochondrial rhythms, including in the *de novo* pyrimidine pathway, before the emergence of long-term protection against DIO. Here we asked whether a timed and transient inhibition of the mitochondrial enzyme DHODH could reproduce a functionally relevant subset of this reprogramming. We show that the short-half-life DHODH inhibitor BAY2402234, administered at ZT0 but not at ZT12, restores mitochondrial oxidative dynamics, increases nighttime carbohydrate use, and, upon repeated dosing, prevents weight gain and reduces hepatic triglycerides in HFD-fed male mice without reducing food intake. The choice of sex is in line with reports that TRF improves metabolic health in both sexes but prevents weight gain only in males^11^. To our knowledge, this is the first demonstration that timed inhibition of a mitochondrial enzyme is sufficient to prevent diet-induced obesity. Notably, although DHODH inhibition did not reproduce the overall metabolite-abundance changes seen with TRF, it did reproduce a time-dependent phenotype. These data indicate that, in this setting, the timing of pathway engagement is a stronger determinant of efficacy than bulk metabolite abundance alone.

Mechanistically, DHODH is unique because it is the only enzyme of the *de novo* pyrimidine pathway located within mitochondria^41,46^. It converts DHO to orotate while reducing CoQ (ubiquinone) to CoQH₂ (ubiquinol)^47^, feeding electrons to complex III. The orchestration of rhythmic respiration is facilitated by the temporal coordination of mitochondrial OXPHOS genes, suggesting that the diurnal compartmentalization of mitochondrial oxidative metabolism is evolutionarily advantageous for optimizing metabolic output^48^.

Uridine, derived from UMP in the *de novo* pyrimidine pathway, regulates energy homeostasis^35^ and exhibits circadian clock-dependent diurnal variation^36^; these rhythms are disrupted by HFD and restored by TRF in the same metabolic context^9,35,37^. In our setting, uridine supplementation only partially reversed the metabolic benefits of BAY2402234, indicating that restoration of pyrimidine/uridine pools is sufficient to blunt part of the phenotype but not to abolish it. Together with the marked difference between ZT0 and ZT12 dosing despite comparable pharmacokinetics, this argues that metabolic context and dosing time override substrate abundance alone. Thus, the efficacy of DHODH inhibition depends less on when pyrimidine intermediates are most abundant than on when the mitochondrial oxidative program is most permissive to brief pathway perturbation.

Notably, although *de novo* pyrimidine intermediates exhibit diurnal oscillations, these rhythms are unlikely to be explained by rhythmic *Dhodh* expression itself. In our study, DHODH protein abundance did not show detectable diurnal rhythmicity in livers of DIO mice under either *ad libitum* feeding or TRF, and published transcriptomic and proteomic resources likewise do not support robust circadian oscillation of *Dhodh* mRNA^11,39,49^ or protein levels^22,50–53^. Together, these observations argue that DHODH expression is not circadian clock-controlled in this context. Rather, its chronopharmacological value appears to arise because a non-rhythmic enzyme is embedded within a rhythmic metabolic network, making the timing of inhibition, rather than the timing of expression, the key determinant of outcome.

In addition to its metabolic effects, timed DHODH inhibition remodeled mitochondrial architecture in a manner similar to short-term TRF. DHODH inhibitors have been reported to increase MFN1/2 expression and promote mitochondrial elongation in cells^31^, and elongated networks are often associated with improved bioenergetic efficiency^54^. Consistent with this framework, ZT0 BAY2402234 normalized the bulbous mitochondrial morphology observed in HFD livers and coincided with functional improvements, supporting the idea that mitochondrial dysfunction in DIO is amenable to chronotherapeutic DHODH inhibition. Because mitochondrial network state can, in turn, influence circadian clock function^27^, improved mitochondrial architecture may also contribute by reinforcing temporal organization of mitochondrial metabolism.

Approved DHODH inhibitors such as leflunomide or teriflunomide have half-lives of ∼2 weeks and are therefore poorly suited for time-of-day–specific inhibition. Moreover, DHODH blockade has generally not been associated with weight loss in those settings. Although Tevonin-1, which inhibits sirtuins and also targets DHODH^33^, prevented weight gain and improved glycemia in HFD-fed rats^55^, the specific contribution of DHODH inhibition could not be isolated. Conversely, a 37-day, once-daily BAY2402234 treatment in *db/db* mice failed to prevent weight gain but decreased food intake through increased GDF15^32^. This study did not test time-of-day effects, and the altered leptin signaling and anorexigenic response in *db/db* mice likely explain the discrepancy with our results. Similarly, the longer half-lived DHODH inhibitor Brequinar produced broader metabolic benefits than BAY2402234 but together with stronger anorexigenic effects. Altogether, these reports are consistent with the idea that in long-exposure or leptin-deficient conditions, the metabolic benefit of DHODH inhibition is mainly food-intake-driven, whereas in our short-pulse, ZT0-restricted paradigm the benefit is time-of-day-driven.

Although these data support DHODH as a rhythm-sensitive metabolic node, several limitations should be acknowledged. First, because all *in vivo* experiments were performed in male mice, these conclusions formally apply to males and should be validated in females. Second, most metabolite measurements relied on whole-liver extracts or isolated mitochondria, approaches that average signals across distinct subcellular pools and may therefore dilute localized or time-dependent changes; future compartment-resolved metabolomics will be required to address this directly. Third, although the time-of-day dependence of BAY2402234 efficacy could in principle reflect differences in tissue exposure or bioavailability, the physicochemical properties, tissue distribution and similar pharmacokinetics observed at ZT0 and ZT12 argue against this as the main explanation. Moreover, the present data do not establish that the anti-obesity effect of timed BAY2402234 administration is solely mediated by hepatic DHODH inhibition. In addition, although our calorimetry and mitochondrial analyses support a time-dependent effect of BAY2402234 on whole-body energy handling, the precise mechanisms underlying energy dissipation remain to be established. Finally, DHODH inhibition has been linked to impaired adipocyte differentiation and mitochondrial function^56^, and TRF benefits require rhythmic adipocyte creatine metabolism^57^; together with the fasting-induced shift of uridine production toward adipose tissue^35^, this raises the possibility that peripheral tissues relay or amplify part of the hepatic effect described here. This possibility will need to be tested directly in future chronopharmacological studies.

In conclusion, we identify a chronopharmacological strategy that targets mitochondrial rhythmic activity. Timed inhibition of DHODH at ZT0 restores rhythms of mitochondrial oxidative functions and improves metabolic adaptation to HFD, independently of changes in feeding schedule, a key distinction from nocturnal TRF. These findings support the broader notion that time-of-day can be used as an additional lever to exploit mitochondrial and pyrimidine pathways in metabolic disease.

## ACKNOWLEDGMENTS

We are grateful to the Genomic Core Facility of Nantes GenoA, member of Biogenouest, for the use of its resources and for its technical support. We are grateful to the SCIAM Department in Angers for help with electron microscopy, especially F. Manero, for technical assistance for electron microscopy imaging. We thank Pr. Chih-Hao Lee and Dr. Ganna Panasyuk for comments.

This research was supported by INSERM ATIP-Avenir (R16067NSRSE17002NSA), the French National Research Agency under the ANR R20109NN and under the “Programme Investissement d’Avenir” (NExT (I-SITE); ANR-16-IDEX-0007), the French Regional Council of Pays de la Loire (2017-02946/02947), “Nantes Métropole”, “Fondation Genavie”, Centre de Recherche en Nutrition Humaine (CRNH) Ouest and the Société Francophone du Diabète (Allocation SFD Jeune Chercheur Francophone 2018).

## Author contributions

1. Conceptualization: D.J. (lead) and F.A.;
2. Data curation : F.A. and D.M.;
3. Formal Analysis : F.A. and D.M.;
4. Funding acquisition : D.J. (lead), F.A. and D.M.;
5. Investigation : F.A. (lead), D.J., M.D., Y.F., I.N., M.C., C.B., S.L.L., E.L.Q., A.C. and D.M.;
6. Methodology : F.A. (lead), D.J., and D.M.;
7. Project administration : D.J.;
8. Supervision : D.J. (lead), and D.M.;
9. Visualization : F.A. (lead), D.J., and D.M.;
10. Writing – original draft : D.J. (lead), F.A. and D.M.;
11. Writing – review & editing : F.A., D.J., M.D., Y.F., I.N., M.C., C.B., S.L.L., E.L.Q., A.C., X.P., C.P. and D.M.

## The authors declare the following competing interests

D.J. reports consulting and advisory roles for Aviitam, Boehringer Ingelheim, AbbVie, AstraZeneca, Novo Nordisk, Eli Lilly, Pfizer, Roche, and Fitforme; investigator roles in phase 3 clinical trials for Eli Lilly, Novo Nordisk, and Roche; and travel reimbursement from Novo Nordisk and Eli Lilly. None of these relationships relate to the present submitted work, and they had no role in the conception, execution, analysis, interpretation, or publication of this study.

## METHODS

### Resource availability Lead Contact

Further information and requests for resources and reagents should be directed to the lead contact, David Jacobi, and will be fulfilled upon reasonable request.

### Materials Availability

This study did not generate new unique reagents.

### Data and code availability statement

RNA-sequencing data have been deposited in the Gene Expression Omnibus under accession GSE230775. This study does not report original code. Any additional information required to reanalyze the data reported in this paper is available from the corresponding authors upon reasonable request.

### Animals

The care and use of animals were approved by the local ethics committee (Comité d’éthique Pays de la Loire, France; project APAFIS#10431) and followed directive 2010/63/EU. Male C57BL/6J mice were housed in ventilated cabinets under a 12 h light/12 h dark cycle (ZT0: lights on; ZT12: lights off), 4–5 mice per cage, with free access to water. Male mice were used to maintain direct comparability with prior HFD/TRF studies, including reports in which TRF prevented weight gain most robustly in males^11^. An overview of the animal experimental designs is provided in Extended Data Table 1.

### TRF experiments

After 4 weeks of acclimation, 9–10-week-old mice were fed an HFD (60% kcal from fat; D12492, Research Diets). Mice were either fed *ad libitum* or restricted to the dark phase (TRF; 8–9 h feeding window) as previously described to maintain isocaloric intake with *ad libitum* controls^9,10^. Whole-body metabolism was assessed by indirect calorimetry (Panlab, Harvard Apparatus) for 3 consecutive days in individually housed mice. GTTs and insulin tolerance tests (ITTs) were performed on fasted mice. For TRF experiments, mice were fasted for 16 h (starting at ZT22) for GTTs or 3 h (starting at ZT13) for ITTs, in accordance with Chaix *et al*.^10^ Mice received 1.5 g/kg body weight glucose (GTT) or 1 U/kg insulin (ITT).

### Genetic *Dhodh* knockdown

*Dhodh* knockdown was induced in mice maintained on standard chow 2 weeks before HFD feeding under *ad libitum* or TRF conditions. Mice were injected retro-orbitally with AAV8-sh-*Dhodh* or control AAV8 (2 × 10¹¹ viral particles/mouse; Vector Biolabs). Livers were collected at ZT6.

### Pharmacological DHODH inhibition

For pharmacological experiments, mice fed an *ad libitum* HFD received intraperitoneal BAY2402234 (1.6 mg/kg; HY-112645, Cliniscience) at ZT0 or ZT12. IP injections were given daily for short experiments (≤4 days) and every other day for longer experiments. Based on pharmacokinetic data showing low residual BAY2402234 levels at 24 h across serum, liver, eWAT and brain, the every-other-day schedule was chosen to avoid drug accumulation during long protocols, as previously done for nobiletin^15^. Mice were euthanized at the indicated time points after the last injection.

### Uridine supplementation

For uridine rescue experiments, uridine (Sigma) was added to the drinking water at 5 g/L to reach ∼400 mg/kg/day. This has been demonstrated to elevate plasma uridine levels^58^.

### Specific protocols

- Food-intake experiment (Extended Data Fig. 7): BAY2402234 (1.6 mg/kg) was injected daily for 7 consecutive days at ZT0 or ZT12; food intake and light/dark distribution were recorded.
- Acute BAY2402234 calorimetry (Fig. 4f): mice were switched to HFD for 3 days, received a single IP injection of BAY2402234 (1.6 mg/kg) at ZT0 or ZT12, and were immediately placed in metabolic cages for 48 h recordings.
- FGF21/GDF15 quantification: blood and liver were collected 4 h and 16 h after the last BAY2402234 injection at ZT0 or ZT12.

### Isolation of liver mitochondria

Liver mitochondria were isolated by differential centrifugation as previously described, with minor modifications ^26^. All manipulations were then performed on ice or at 4°C. Briefly, ∼150 mg of liver was washed twice, minced, and homogenized in mitochondrial isolation buffer (MIB: 70 mM sucrose, 210 mM mannitol, 5 mM HEPES, 1 mM EGTA, and 0.5% BSA fatty acid free). Samples were centrifuged at 800 g for 8 min and mitochondria-containing supernatants filtered through a 70 µm filter. The filtered supernatants were centrifuged for 8 min at 8.000 g. The resulting pellet was rinsed and centrifuged 5 min at 8000 g. Mitochondrial pellets were suspended in 100 µL of MIB. If the mitochondrial suspension was not used directly, mitochondria were pelleted and stored at -80°C.

### Bioenergetics analyses

The oxygen consumption rate was determined on a Seahorse XF HS Mini Analyzer. For mitochondrial coupling assays, isolated mitochondria were incubated in a mitochondrial assay buffer (70 mM sucrose, 220 mM mannitol, 10 mM KH_2_PO_4_, 5 mM MgCl_2_, 2 mM HEPES, 1 mM EGTA, and 0.2% fatty acid free BSA, pH 7.2) with 10 mM succinate and 2 μM rotenone.

### Immunoblots

Liver homogenates and mitochondrial extracts were prepared from frozen liver and from isolated mitochondrial pellets, respectively, in lysis buffer containing 8 M urea, protease inhibitor (cOmplete™, EDTA-free; Roche), phosphatase inhibitor (PhosSTOP; Roche) and sirtuin inhibitors (AGK7 and salermide, Bertin) at the manufacturer’s recommended concentrations. Proteins were separated by SDS–PAGE on Novex gels (Thermo Fisher Scientific) and transferred to PVDF membranes (Trans-Blot; Bio-Rad). Membranes were blocked and incubated with the following primary antibodies: Total OXPHOS cocktail (Invitrogen, 458099), FGF21 (Abcam, ab171941), β-actin (Cell Signaling Technology, 13E5; #4970S) and Histone H4 (Thermo Fisher Scientific, 16047-1-AP). Signals were acquired on a Bio-Rad imaging system and quantified in ImageJ; band intensities were normalized to total protein as assessed by Naphthol blue black total-protein staining (Sigma).

### Metabolomic analyses of liver and hepatic mitochondria

Targeted metabolomics of small polar metabolites in central carbon metabolism was performed using previously described extraction and LC–MS procedures^59^. Liver and mitochondrial samples were processed with the same extraction solvent (50% methanol, 30% acetonitrile, 20% water), as in Mackay *et al.*^59^. Quality assurance and quality control followed the recommendations of the Metabolomics Quality Assurance & Quality Control Consortium (mQACC)^60^, including instrument maintenance (Thermo), SOPs for extraction and storage, pooled inter-study QC samples, process and extraction blanks, system-stability blanks, a long-term interlaboratory reference mix, and randomized blinded samples; no significant drift was detected across QC samples.

For liver, tissue was flash-frozen in liquid N₂ and pulverized; 40 mg were extracted in 1 mL of solvent. For hepatic mitochondria, isolated pellets were resuspended in KPBS to 250 µg total protein in 35 µL and extracted in 250 µL of the same solvent. All samples were vortexed for 5 min at 4°C and centrifuged at 16,000 g for 15 min at 4°C; supernatants were collected and stored at –80°C until analysis.

LC–MS was carried out on a Q Exactive Plus Orbitrap mass spectrometer equipped with an Ion Max source and HESI II probe, coupled to a Dionex UltiMate 3000 UPLC (Thermo). Five microlitres of extract were injected onto a ZIC-pHILIC column (150 × 2.1 mm, 5 µm) with matching guard column (20 × 2.1 mm, 5 µm) (Millipore). Mobile phase A was 20 mM ammonium carbonate, 0.1% ammonium hydroxide (pH 9.2); mobile phase B was 100% acetonitrile. The gradient (200 µL/min) was: 0–20 min, 80%→20% B; 20–20.5 min, 20%→80% B; 20.5–28 min, 80% B (re-equilibration). The MS was operated in polarity-switching full-scan mode (m/z 75–1,000), resolution 35,000 at m/z 200, AGC target 1×10⁶, max IT 250 ms; spray voltage 2.5 kV, capillary 320°C, sheath gas 20, auxiliary gas 5. Raw data were acquired in Xcalibur (Thermo) and processed in TraceFinder (v3.3.350, Thermo). Metabolites were identified by exact m/z and by matching retention time to an in-house library built from LSMLS™ / IROA Technologies standards; peak areas were exported for downstream statistical and rhythmicity analyses.

### Quantitative metabolomic analyses of serum and tissue

Quantitative targeted metabolomics analyses were performed by liquid chromatography-tandem mass spectrometry (LC-MS/MS) consisting of a Xevo^®^ TQD mass spectrometer with an ESI interface coupled to Acquity H-Class^®^ UPLC^TM^ device (Waters Corporation, Milford, MA, USA). Serum samples did not require any preparation prior to extraction. On the other hand, tissue samples (liver, adipose tissue, and brain) were pulverized, weighed and diluted 1:5 in PBS (5 mL per gram of sample). Homogenization of the samples was then performed mechanically using small metal balls and a tissue lyser (TissueLyser II, Qiagen). A portion of the recovered lysate was diluted 1:5 in UPLC ultrapure water before proceeding with a protein assay (Thermofisher, Pierce BCA Protein Assay Kit), to normalize the concentrations obtained by LC-MS/MS by the total protein concentration.

### Uridine, dihydro-orotate (DHO) and orotate measurements

To quantify uridine, dihydro-orotate (DHO) and orotate, 1 mM stock solutions of DHO (Sigma, D7003), orotate (Sigma, O2750) and uridine (Sigma, U3750) were prepared in UPLC-grade water and serially diluted to generate calibration standards from 0 to 100 µM in the corresponding matrix. 5-Fluoroorotate (Sigma, F5013) at 50 µM was used as an exogenous internal standard to correct for extraction efficiency and matrix effects.

#### Serum

Ten microlitres of serum were mixed with 10 µL of internal standard (50 µM) and 80 µL of cold acetonitrile, vortexed (10 s) and centrifuged (10,000 g, 10 min, 10°C). The supernatant (∼80 µL) was transferred to LC–MS vials.

#### Liver and adipose tissue

Tissue homogenates (100 µL) were spiked with 25 µL of internal standard (50 µM), extracted with 400 µL of cold acetonitrile, vortexed (10 s) and centrifuged (10,000 g, 10 min, 10°C). The supernatant (≈450 µL) was collected, dried under nitrogen and reconstituted in 80 µL of acetonitrile. Calibration samples were processed in parallel following the same procedure.

Five microlitres of each extract were injected onto a HILIC BEH amide column (1.7 µm, 2.1 × 100 mm; Waters) kept at 45°C. Metabolites were separated at 0.8 mL/min with mobile phase A (10 mM ammonium acetate, 0.1% formic acid) and mobile phase B (98% acetonitrile, 0.1% formic acid) using the following program: 0–0.5 min, 1% A; 0.5–4.5 min, 1%→45% A; 4.5–5.0 min, 45% A; 5.0–5.5 min, return to 1% A; 5.5–7.0 min, re-equilibration. Detection was performed by LC–MS/MS with ESI in negative mode (capillary 3.5 kV; source 130°C; desolvation 350°C, 600 L/h). MRM transitions were: DHO m/z 156.9→112.9 (cone 25 V, collision 8 V), orotate m/z 154.9→110.9 (cone 20 V, collision 10 V), uridine m/z 245.0→112.9 (cone 15 V, collision 10 V), and 5-fluoroorotic acid m/z 172.9→128.9 (cone 15 V, collision 8 V). Data were acquired and processed with MassLynx and TargetLynx (v4.1, Waters), and metabolite concentrations were calculated from matrix-matched calibration curves.

### Ubiquinone redox status

The determination of the redox status of CoenzymeQ9 (CoQ9) involves the detection of its oxidized form, CoQ9 (ubiquinone), and its reduced form CoQ9H_2_ (ubiquinol). The concentration of each form was determined using standard solutions. CoQ9 (Bertin, ref. 16866) and ubiquinol (Sigma, ref. 1705334) standards (0-50 µM) were prepared at 1 mM in methanol. A 100 µL aliquot of each standard solution was taken for extraction. The internal standard CoQ_4_ (Sigma, ref. C2470), an exogenous form in rodents, was prepared in methanol at 10 µM. To extract CoQ from liver, 20 µL of internal standard was added to 100 µL of tissue homogenate or standard solutions, and then 900 µL of a methanol/chloroform mixture (2:1, v:v) was added. After 10 seconds of vortexing, the samples were centrifuged at 10,000 g for 10 min (10°C). The supernatant (800 µL) was collected and transferred to a glass vial compatible with LC-MS/MS and dried under a gentle stream of nitrogen. Dried samples were solubilized in a mixture of 50% acetonitrile in water containing 0.1% formic acid.

Samples (10 μL) were injected onto a BEH-C18 column (i.d. 1.7 μm; 2.1 × 50 mm; Waters Corporation) held at 60°C, and compounds were separated with a linear gradient of mobile phase B (50% acetonitrile, 50% isopropanol 0.1% formic acid, 10 mM ammonium formate) in mobile phase A (5% acetonitrile, 10 mM ammonium acetate, 0.1% formic acid) at a flow rate of 400 μL/min. Mobile phase B was kept constant for 0.5 min at 40%, linearly increased from 40% to 95% for 4 min, kept constant for 2 min, returned to the initial condition over 0.5 min, and kept constant for 0.5 min before the next injection. Targeted compounds were then detected by the mass spectrometer with the ESI interface operating in the positive ion mode (capillary voltage, 3.0 kV; desolvatation gas (N_2_) flow and temperature, 1000 L/h and 400°C; source temperature, 120°C). The multiple reaction monitoring mode was applied for MS/MS detection: CoQ9, m/z 812.6 → 197.1 (cone and collision voltages set at 25 and 20 V, respectively); CoQ9H_2_, m/z 814.6 → 197.1 (cone and collision voltages set at 30 and 25 V, respectively); CoQ_10_, m/z 880.7 → 197.1 (cone and collision voltages set at 30 and 25 V, respectively); CoQ_10_H_2_, m/z 882.7 → 197.1 (cone and collision voltages set at 30 and 25 V, respectively); CoQ_4_, m/z 455.3 → 197.0 (cone and collision voltages set at 30 and 20 V, respectively). Data acquisition and analysis were performed using MassLynx^®^ and TargetLynx^®^ software, respectively (version 4.1; Waters). Compound concentrations were calculated using calibration curves plotted from standard solutions.

### Pharmacokinetics of BAY2402234

Male C57BL/6J mice fed an HFD were injected intraperitoneally with BAY2402234 (1.6 mg/kg) at either ZT0 or ZT12 (n = 4 per time point and condition). Blood (serum) and tissues (liver, epididymal white adipose tissue and brain) were collected at 0.5, 2, 4, 8 and 24 h after injection and immediately snap-frozen in liquid N₂. For quantification, serum and tissue homogenates were extracted with acetonitrile (1:5, v/v), centrifuged (10,000 g, 10 min, 10°C), and the supernatants were dried under N₂ and reconstituted in 100 μL of 25% acetonitrile / 0.1% formic acid. Calibration curves (0–5 μM) were prepared by spiking BAY2402234 into matched serum or tissue lysates and processed in parallel.

Ten microlitres of each sample were injected on a BEH C18 column (1.7 μm, 2.1 × 50 mm; Waters) kept at 60°C. Analytes were eluted at 400 μL/min with mobile phase A (5% acetonitrile, 0.1% formic acid) and mobile phase B (100% acetonitrile, 0.1% formic acid) using the following program: 0–0.5 min, 5% B; 0.5–4.0 min, 5%→95% B; 4.0–4.5 min, 95% B; 4.5–5.0 min, re-equilibration at 5% B. BAY2402234 was detected by LC–MS/MS on a triple-quadrupole instrument equipped with an ESI source operated in positive mode (capillary 3.0 kV; source 120°C; desolvation 400°C, 1000 L/h). Multiple-reaction monitoring (MRM) was used with the transition m/z 521.0 → 376.0 (cone 50 V, collision 28 V). Data were acquired and processed with MassLynx and TargetLynx (v4.1, Waters). Concentrations were obtained from the external calibration curves, and PK parameters (including AUC) were calculated by non-compartmental analysis.

### 3’UTR sequencing and analysis

RNA extraction was performed on frozen liver using Nucleozol (Macherey Nagel). The 3’seq RNA profiling protocol was performed as previously described^61^. Libraries were prepared from 10 ng of total RNA in 4 µl. Poly(A) tails of mRNAs are labeled with universal adapters, well-specific barcodes, and unique molecular identifiers (UMIs) during template-switching reverse transcriptase. Barcoded cDNAs from multiple samples are then pooled, amplified, and labeled using a transposon fragmentation approach that enriches the 3′ ends of the cDNA: 200 ng of full-length cDNA are used as input to the Nextera™ DNA Flex Library Prep kit (ref #20018704, Illumina) and Nextera™ DNA CD indexes (24 indexes, 24 samples) (ref #20018707, Illumina) according to the manufacturer’s protocol (Nextera DNA Flex Library Document, ref #1000000025416 v04, Illumina). The library size is controlled on the 2200 Tape Station Sytem (Agilent Technologies). A 350-800 bp long library is run on a NovaSeq 6000 using the NovaSeq 6000 SP Reagent Kit 100 cycles (ref #20027464, Illumina) with reads of 17*-8-105* cycles.

Primary analysis was performed as previously described^62^. For analysis, the raw fastq pairs correspond to the following criteria: the 16 bases of the first read correspond to 6 bases for a barcode specific to the designed sample and 10 bases for a unique molecular identifier (UMI). The second read (104 bases) corresponds to the sequence of the captured poly(A) RNAs. Demultiplexing was performed with an in-house python script. Raw paired-end fastq files were transformed into a single-end fastq file for each sample.

Alignment on Ensembl *Mus Musculus* annotation (assembly GRCm38/mm10) reference transcriptome^63^ was performed using bwa^64^. Aligned reads were parsed and UMIs counted for each gene to create an expression matrix containing the absolute abundance of mRNAs in all samples. Reads aligned on multiple genes or containing more than three mismatches with the reference were discarded. The expression matrix was normalized using the R package of DESeq2^65^.

### Rhythmicity analysis

Analysis of both mRNA sequencing and metabolomics data were performed with CircaCompare^38^. Resulting *p-values* were adjusted using the Benjamini-Hochberg method to control for the false-discovery rate (FDR). Unless otherwise indicated, genes expression or metabolites abundances with FDR < 0.05 were considered as rhythmic.

### Functional annotation (GO TERM)

Gene ontology (GO) analysis was performed with Pantherdb^66^. Enrichment test was performed for Cellular component (CC_all). Top 10 enriched categories with adjusted p-values (Benjamini–Hochberg) smaller than 0.05 were reported.

### Joint analyses of transcriptome and metabolome data

Integrated metabolic pathway analyses were performed by combining metabolomics and gene expression data using Metaboanalyst 5.0^67^. Topology measure was set to “degree centrality” and enrichment test in metabolic pathways was performed using hypergeometric tests while tight integration was ensured by combining queries.

### Transmission electron microscopy

Electron microscopy was conducted at the SCIAM Department in Angers, France. Following heart perfusion of 15 mL 2% paraformaldehyde, 2.5% glutaraldehyde in 0.1 M Na cacodylate HCl buffer (pH 7.3), small liver blocks were excised from the left lobe and placed in the same fixative at 4°C for 24 h. The blocks were postfixed with 1% OsO_4_/1.5% KFeCN_6_ for 2 h, washed in water, and incubated in 1% aqueous uranyl acetate for 1 h, followed by dehydration through 50%, 70%, 90%, and 100% ethanol. Samples were embedded in TAAB and polymerized at 60°C for 48 h. Ultrathin sections (60 nm) were cut on a Reichert Ultracut-S microtome, stained with lead citrate and examined in a JEM1400, JEOL camera Transmission electron microscope. Images were recorded with an ORIUS SC1000 11 MP camera. To determine the location of hepatocytes relative to the liver lobules, additional sections were cut at 0.5 μm and stained with toluidine blue. Photographs were taken at 5000X magnification for 8 fields/liver, 70-80 μm from the centrilobular vein. Individual mitochondria were manually delineated using ImageJ software for determination of mitochondrial surface, density (number of mitochondria/surface area), and coverage (total mitochondrial surface/total cellular surface area). Mitochondria mitochondrial surface area was measured and untruncated mitochondria were retained for the analysis. For each mouse, the total liver cell surface evaluated was ∼1,500 μm^2^ (8 fields of 187.8 μm^2^ each).

### qPCR

Relative expression was determined by SYBR Green-based real-time PCR using *Gapdh* as an internal standard. Gene expression was analyzed with standard curve method^68^.

### Serum GDF15 quantification

Serum GDF15 was measured with Mouse/Rat GDF15 Quantikine ELISA Kit (R&D Systems), following the manufacturer recommendations.

### Liver triglyceride content

Liver triglyceride concentration was determined with the Infinity™ Triglycerides Liquid Stable kit (Thermo Fisher Scientific) based on an enzymatic colorimetric reaction.

### Statistical Tests

Two-parameter analyses for samples from *in vivo* studies were determined using Welch’s t-test. Statistics for multiparameter analyses were determined by one-way or two-way ANOVA followed by Bonferroni post hoc tests. Rhythmicity analyses were performed using CircaCompare version 0.2.0.^38^ Unless otherwise indicated, *p* <0.05 is considered as significant.

### Data analysis of BAY/VEH injection at ZT0/12 in the long term

A linear model was fitted and applied to the normalized log2-transformed metabolites levels. We used the anova function to compare a full model (with the explanatory variables) and reduced model with only an intercept to identify metabolites that are significantly different across groups (BAY injected at ZT0 and sampled at ZT16 *i.e.* BAY_ZT00_16 or BAY_ZT00_4 or VEH_ZT00_16 or VEH_ZT00_4). Subsequently, we contrast all possible combination of groups within the linear model. Finally, *p-values* of the individual contrasts were used in the confirmation stage. *P-values* were corrected for multiple testing using stageR^69^ with the ANOVA result used during the screening stage, while *p-values* of the individual contrasts were used in the confirmation stage.

## Excel-format tables

**Supplementary Data 1 (related to Fig. 1):** log_2_ median-normalized liver and mitochondrial metabolite levels in short- and long-term TRF with downstream differential rhythmicity analyses.

**Datasheet1:** differential rhythmicity analysis of AL versus TRF in Short-term in all extracts using the Circacompare function of the CircaCompare package.

**Datasheet2:** differential rhythmicity analysis of AL versus TRF in Long-term in all extracts using the Circacompare function of the CircaCompare package.

**Datasheet3**: differential rhythmicity analysis of the overlapping metabolites in mito versus liver extracts in all diet conditions and diet duration (Short and Long term) using the Circacompare function of the CircaCompare package.

**Supplementary Data 2 (related to Extended Data Fig. 2):** log_2_ normalized transcriptomic in short-and long-term TRF and downstream differential rhythmicity and GOTERM enrichment analyses

**Datasheet1**: Differential rhythmicity analysis of AL versus TRF in Short-term using the Circacompare function of the CircaCompare package.

**Datasheet2**: Differential rhythmicity analysis of AL versus TRF in Long-term using the Circacompare function of the CircaCompare package.

**Datasheet3**: GOTERM analyses of genes with rhythmicity imprinted by TRF in Short term.

**Datasheet4**: GOTERM analyses of genes with rhythmicity imprinted by TRF in Long term.

**Supplementary Data 3 (related to Fig. 2):** Integrative pathway enrichment analysis of rhythmic transcript and metabolites.

**Datasheet1:** integrative pathway enrichment analysis of rhythmic transcript and metabolites in short-term TRF.

**Datasheet2:** integrative pathway enrichment analysis of rhythmic transcript and metabolites in long-term TRF.

**Supplementary Data 4 (related to Fig. 4):** log_2_ median-normalized metabolite and rhythmicity analysis in BAY2402234 ZT0 or ZT12 controls.

**Datasheet1:** Differential rhythmicity analysis of DMSO versus BAY injected at ZT00 using the Circacompare function of the CircaCompare package.

**Datasheet2:** Differential rhythmicity analysis of DMSO versus BAY injected at ZT12 using the Circacompare function of the CircaCompare package.

**Supplementary Data 5 (related to Fig. 5):** log_2_ median-normalized mitochondrial metabolome with time-dependent change in BAY2402234 ZT0 or ZT12 and controls

**Datasheet1:** Differential analysis of DMSO versus BAY injected at ZT00; liver samples collected at ZT4 and ZT16 after 4 weeks of treatment; p-values were corrected for multiple testing using stageR.

**Datasheet2:** Differential analysis of DMSO versus BAY injected at ZT00; mitochondria samples collected at ZT4 and ZT16 after 4 weeks of treatment; p-values were corrected for multiple testing using stageR.

**Datasheet3:** Differential analysis of DMSO versus BAY injected at ZT12; liver samples collected at ZT4 and ZT16 after 4 weeks of treatment; p-values were corrected for multiple testing using stageR.

**Datasheet4:** Differential analysis of DMSO versus BAY injected at ZT12; mitochondria samples collected at ZT4 and ZT16 after 4 weeks of treatment; p-values were corrected for multiple testing using stageR.

---

**Extended Data Figure 1 (related to Fig. 1):**
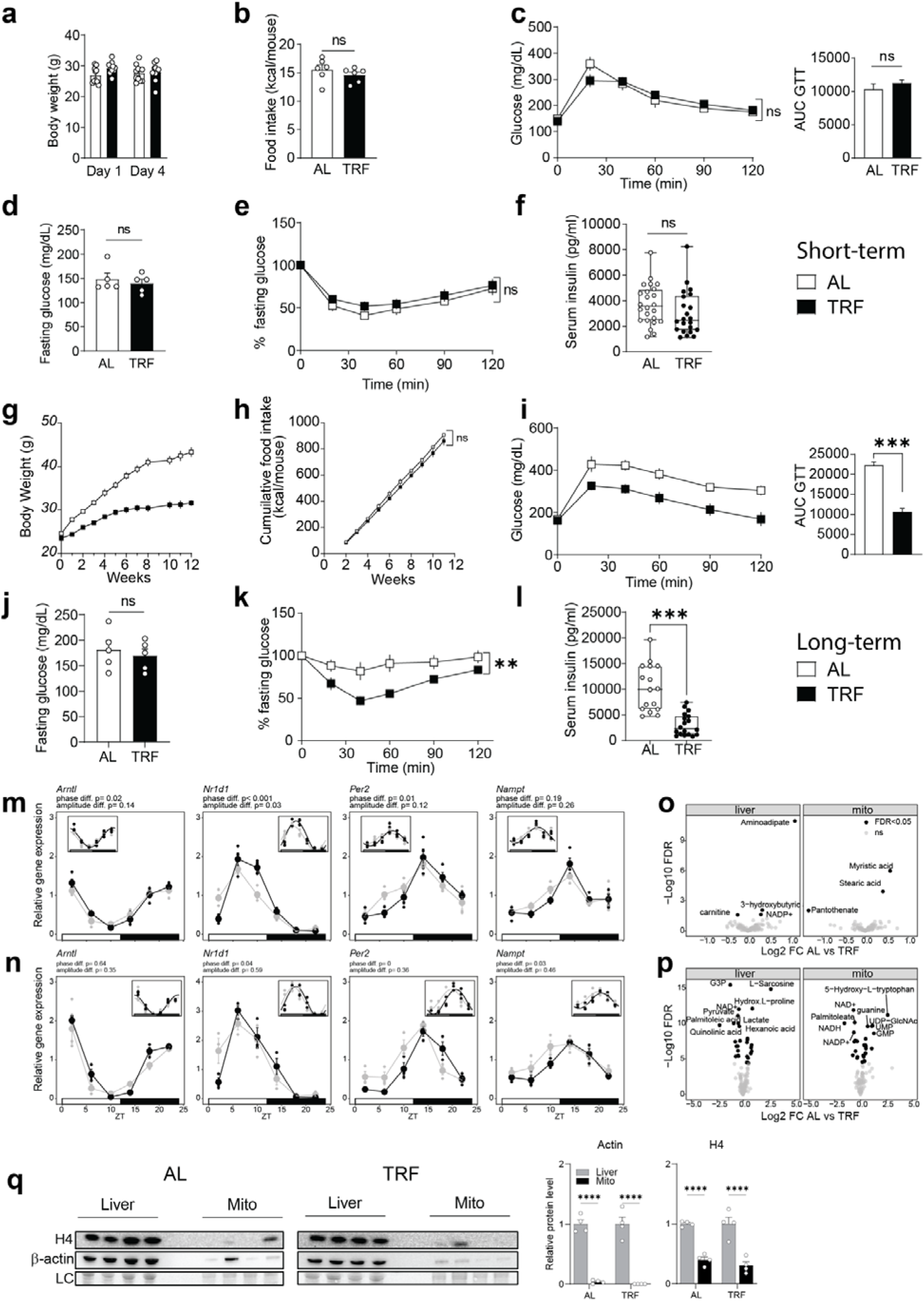
Early promotion of circadian and metabolomic rhythms by TRF. Metabolic phenotyping was performed after short-term (a–f; 4 days, n=24/group) and long-term TRF (g–l; 12 weeks, n=18/group). **(a,g)** Body weight (short-term: start versus end, n=24/group; long-term: trajectory/endpoint, n=18/group). **(b,h)** Cumulative food intake (short-term,n=6/group;long-term,n=5/group). **(c,i)** Glucose tolerance test (GTT, left) and AUC above baseline (right).n=8/group **(d,j)** Fasting glycaemia.n=8/group **(e,k)** Insulin tolerance test (ITT). n=8/group **(f,l)** Serum insulin (short-term, n=24/group; long-term, n=18/group). **(m,n)** Hepatic circadian clock and clock-controlled gene expression across the light/dark cycle (short-term (**m**, n=4/timepoint/group) and long-term TRF (**n**, n=3/timepoint/group)); solid lines indicate CircaCompare fits when significant. **(o,p)** TRF-imprinted changes in liver (left) and mitochondrial (right) metabolite abundance using rhythm-adjusted mean (mesor) comparisons (ANOVA, FDR < 0.05; n as in Fig. 1a design). **(q)** Immunoblot validation of mitochondrial fraction enrichment (β-actin, histone H4 depletion; quantification at right). Loading control (LC): Naphtol Blue Black. n=4/group. Data are mean ±s.e.m. ; **P<0.01, ***P<0.001, ****P<0.0001. All experiments were performed in male C57BL/6J mice.

**Extended Data Fig. 2. Related to Fig. 2.**
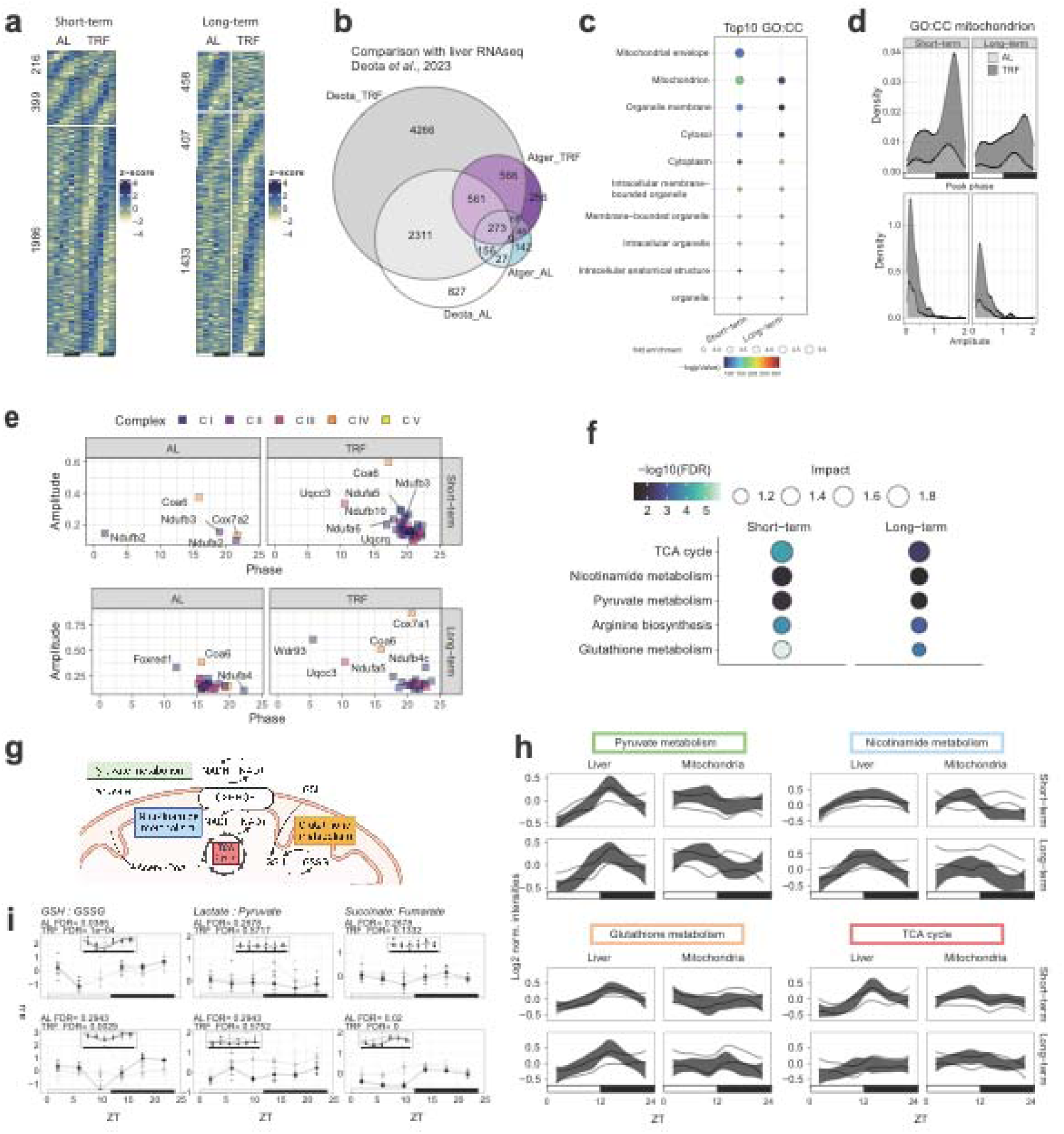
TRF remodels daily mitochondrial oxidative activities. **a,** Number of rhythmic liver transcripts detected by CircaCompare (FDR < 0.05) in short- and long-term TRF. **b,** Overlap of rhythmic transcripts in long-term TRF with Deota et al. 2023. **c,** GO:CC enrichment for TRF-induced rhythmic transcripts. **d,** Peak-phase distribution and amplitude changes under TRF among mitochondria-annotated rhythmic transcripts. **e,** Temporal expression of representative nuclear-encoded mitochondrial respiration genes with TRF-promoted oscillations (n = 3–4 mice per ZT per condition; lines show CircaCompare fit where significant). **f,** Integrative analysis of rhythmic transcriptome with rhythmic liver and mitochondrial metabolomes (MetaboAnalyst) showing enriched oxidative/energy pathways (FDR < 0.05). **g,** Schematic summary of enriched mitochondrial pathways. **h,** Rhythmic metabolite patterns in glycolysis/pyruvate and related pathways supporting mitochondrial redox and acetyl-CoA supply. **i,** Redox-related proxies/ratios (including GSH:GSSG, succinate:fumarate, lactate:pyruvate), and/or TCA/nicotinamide rhythmicity readouts (mean ± s.e.m.). All experiments were performed in male C57BL/6J mice.

**Extended Data Figure 3 (related to Fig. 3):**
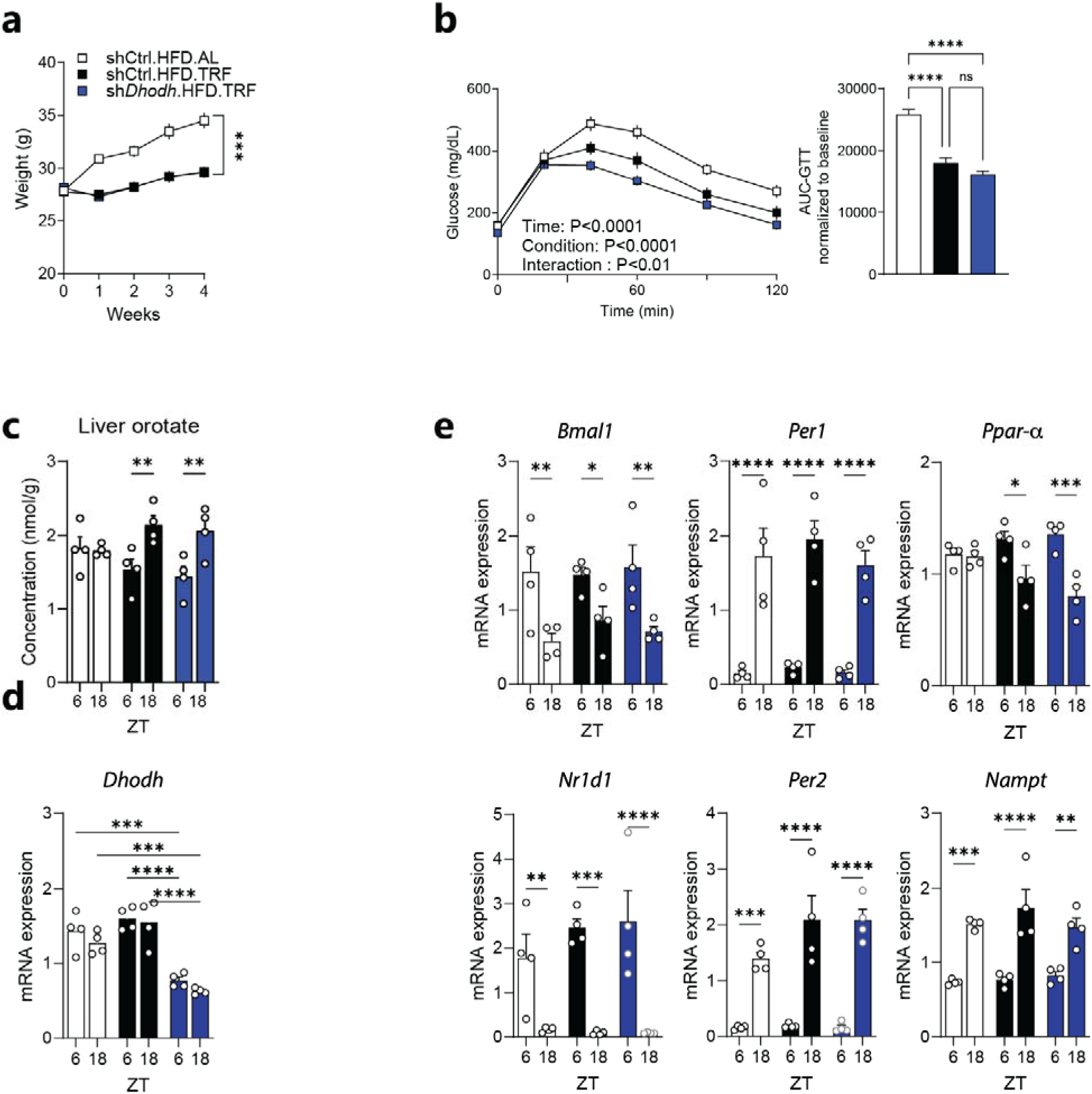
Liver *Dhodh* knockdown under HFD and TRF. **a,** Body-weight evolution over 4 weeks in mice fed a high-fat diet (HFD) ad libitum (sh-Ctrl.HFD.AL) or under time-restricted feeding (sh-Ctrl.HFD.TRF), and in TRF mice with liver *Dhodh* knockdown (sh-*Dhodh*.HFD.TRF). n=8/group. **b,** Glucose tolerance test (GTT) and corresponding AUC (normalized to baseline) in the three groups; n=8/group; P values for time, condition and time × condition interaction are indicated on the plot. **c,** Hepatic orotate concentration at ZT6 and ZT18. n=8/group. **d,** Hepatic *Dhodh* mRNA abundance at ZT6 and ZT18 confirming knockdown efficiency. n=8/group. **e,** RT–qPCR analysis of hepatic clock/metabolic gene expression (*Bmal1, Per1, Per2, Nr1d1* (Rev-erbα), *Ppar*α*, Nampt*) at ZT6 and ZT18 across groups. Data are mean ± s.e.m.; each dot represents one biologically independent mouse (n=8/group). Statistical significance is shown as indicated (\**P* < 0.05, \*\**P* < 0.01, \*\*\**P* < 0.001, \*\*\*\**P* < 0.0001; ns, not significant). All experiments were performed in male C57BL/6J mice.

**Extended Data Figure 4:**
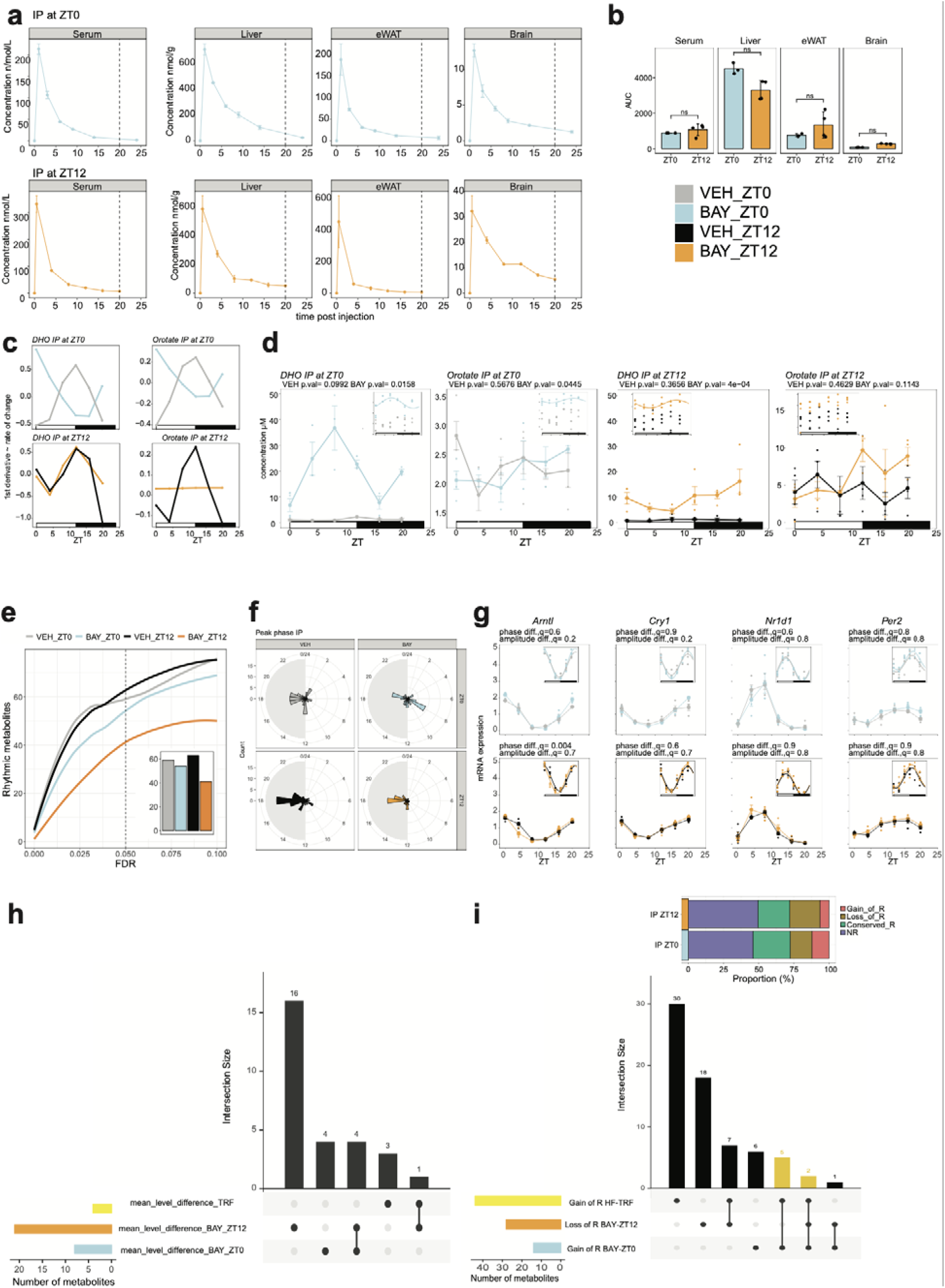
Timing of BAY2402234 administration shapes biodistribution and rhythmic liver metabolism without altering core clock gene expression. **a,** Pharmacokinetics of BAY2402234 in serum, liver, epididymal white adipose tissue (eWAT) and brain after a single intraperitoneal (IP) injection at ZT0 (top) or ZT12 (bottom); tissues collected 0.5, 2, 4, 8 and 24 h post-injection (n = 4 per time point and condition). **b,** Temporal biodistribution of BAY2402234 quantified as area under the curve (AUC) in serum, liver, eWAT and brain for ZT0 versus ZT12 injection. **c,** First-derivative (standardized rate-of-change) plots for hepatic DHO and orotate corresponding to (Fig.4b). **d,** Serum dihydroorotate (DHO) and orotate profiles across the light/dark cycle in mice treated for 4 days with vehicle (VEH) or BAY2402234 administered daily at ZT0 or ZT12 (n = 3–4 per ZT per condition). **e,** Number of rhythmic liver metabolites detected by CircaCompare across false discovery rate (FDR) thresholds (inset, number at 0.05 FDR cutoff). **f,** Peak phase distribution of rhythmic metabolites across conditions (VEH versus BAY; ZT0 versus ZT12 dosing). **g,** Liver RT–qPCR time courses of core circadian clock genes; solid lines indicate CircaCompare fits (P < 0.05) across the light/dark cycle (n = 3–4 per ZT per condition). **h, i,** UpSet plots summarizing overlap between metabolite sets defined by mean-level (mesor) differences (**h**) or rhythmicity changes (**i**) across BAY dosing-time conditions and TRF-associated signatures. Data are mean ± s.e.m. unless indicated. All experiments were performed in male C57BL/6J mice.

**Extended Data Figure 5 (related to Fig. 5):**
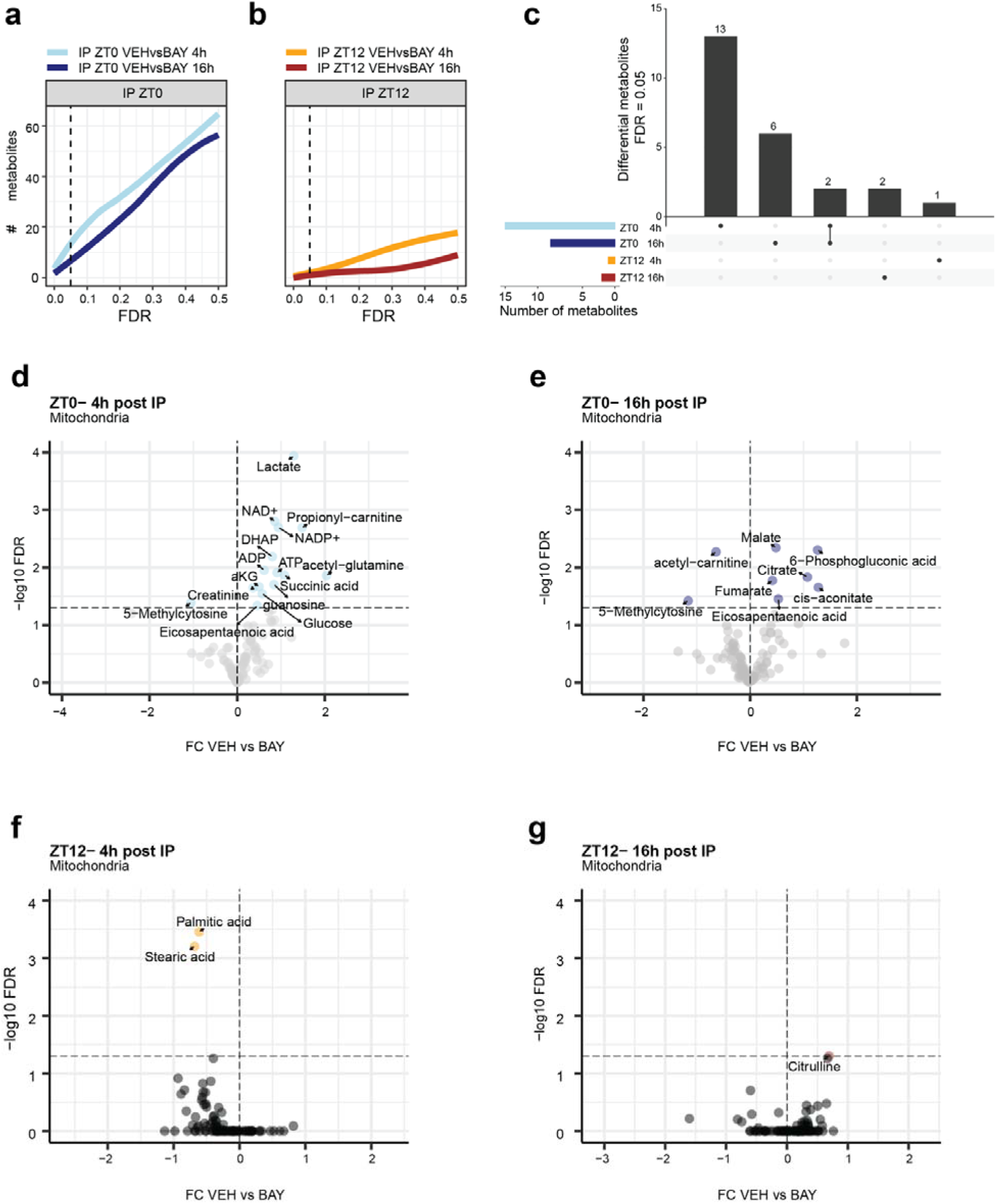
Time- and dosing-time–dependent remodeling of the hepatic mitochondrial metabolome after transient DHODH inhibition. **a,b,** Cumulative number of differential mitochondrial metabolites between vehicle (VEH) and BAY2402234 across false discovery rate (FDR) thresholds, measured 4 h or 16 h after the final intraperitoneal (IP) injection administered at ZT0 (a) or ZT12 **(b)**. **c,** UpSet plot showing overlap of BAY2402234-regulated mitochondrial metabolites across the four conditions (ZT0–4 h, ZT0–16 h, ZT12–4 h, ZT12–16 h) at the indicated FDR cutoff. **d,e,** Differential mitochondrial metabolites after BAY2402234 dosing at ZT0, assessed 4 h **(d)** or 16 h **(e)** post-IP (volcano plots; x-axis, log2 fold change BAY/VEH; y-axis, −log10 FDR); selected bioenergetics and TCA-cycle intermediates are annotated. **f,g,** Same analysis for BAY2402234 dosing at ZT12, 4 h **(f)** or 16 h **(g)** post-IP, showing minimal deviation from VEH. Mitochondria were purified from livers of mice maintained on 12-week HFD and treated during the final 4 weeks with every-other-day IP injections of BAY2402234 or VEH at ZT0 or ZT12; animals were sacrificed at ZT4 or ZT16 after the last injection (n = 4 mice per group). All experiments were performed in male C57BL/6J mice.

**Extended Data Figure 6:**
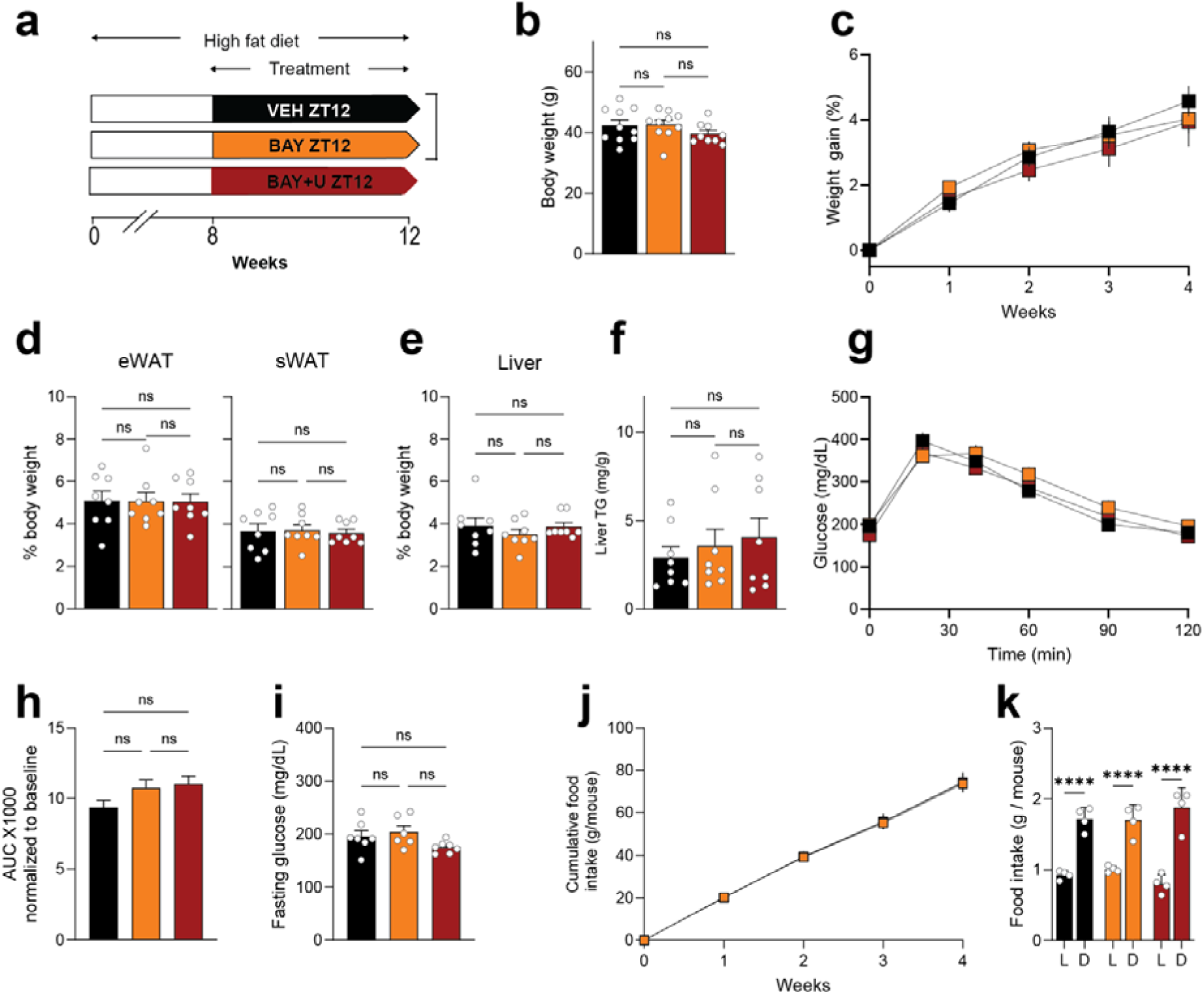
BAY2402234 dosing at ZT12 does not improve metabolic outcomes in diet-induced obese mice, irrespective of uridine supplementation. **a,** Experimental design. Mice were fed a high-fat diet (HFD) for 12 weeks. From week 8 to week 12, mice received intraperitoneal (IP) injections every other day at ZT12 of vehicle (VEH), BAY2402234 (BAY; 1.6 mg/kg), or BAY2402234 together with uridine in the drinking water (BAY+U; 5 g/L) (n = 10/group). **b,** Terminal body weight. n=8/group. **c,** Body-weight gain (%) during the 4-week treatment period. n=8/group. **d,** Epididymal (eWAT) and subcutaneous (sWAT) fat mass expressed as % body weight. n=8/group. **e,** Liver weight as % body weight. n=8/group. **f,** Hepatic triglyceride (TG) content. n=8/group. **g,** Glucose tolerance test (GTT) at week 3, 6 h after the last injection; mice were fasted for 6 h before the test. n=6-7/group. **h,** GTT area under the curve (AUC) corresponding to panel **g**. **i,** Fasting blood glucose. n=7/group. **j,** Cumulative food intake per mouse over the 4-week treatment. n=8-9/group. **k,** Food intake during the light (L) and dark (D) phases (3 independent 24-h measurements per group over the 4 weeks). n=8/group. Data are mean ± s.e.m. ns, not significant; *P < 0.05; **P < 0.01; ***P < 0.001; ****P < 0.0001. One-way ANOVA (or Welch’s t-test, as appropriate) was used for panels b, d–f, h–k; two-way ANOVA with Bonferroni’s post-hoc test was used for panels c and g. All experiments were performed in male C57BL/6J mice.

**Extended Data Figure 7:**
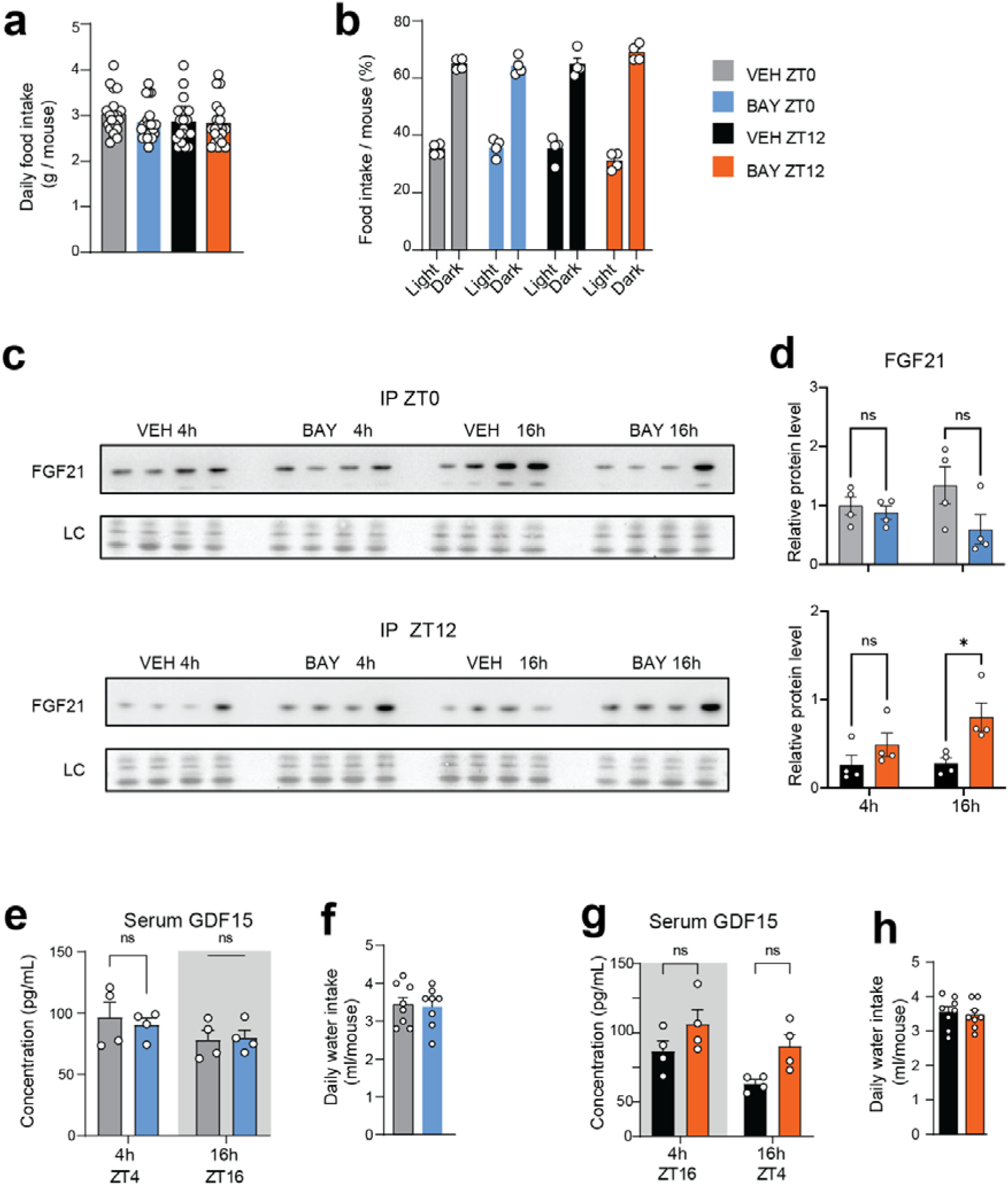
DHODH inhibition does not markedly change feeding or hydration, but modulates FGF21 in a time- and dosing-time–dependent manner. **a,** Daily food intake (g per mouse) measured in HFD-fed mice receiving daily intraperitoneal injections of vehicle (VEH) or BAY2402234 (BAY) at ZT0 or ZT12 for 7 consecutive days (n=20/group). **b,** Distribution of food intake between light and dark phases for the same conditions as in (**a)** (n=4 cages/group). **c,** Representative immunoblots of hepatic FGF21 collected 4 h and 16 h after VEH or BAY injection at ZT0 (top) or ZT12 (bottom). LC, loading control. n=4/group/timepoint. **d,** Densitometric quantification of FGF21 normalized to LC at 4 h and 16 h after injection; ns, not significant; *P* < 0.05. **e,g,** Serum GDF15 measured 4 h and 16 h after VEH or BAY injection at ZT0 (**e**; sampling at ZT4 and ZT16) or ZT12 (**g**; sampling at ZT16 and ZT4), respectively. n=4/group/timepoint. **f,h,** Daily water intake following injections at ZT0 (**f**) or ZT12 (**h**). n=8/group/timepoint. Data are mean ± s.e.m. Statistical comparisons were performed between VEH and BAY at each matched time point; ns, not significant; \**P* < 0.05. All experiments were performed in male C57BL/6J mice.

**Extended Data Table 1:** Design of animal experiments and related figures and supplementary data.

| Experiment ID | Experiment | Diet | Duration | BAY2402234 Dosing schedule | Comparison | Related figures | Related Supplementary Data |
| --- | --- | --- | --- | --- | --- | --- | --- |
| Exp1 | Short-term time-restricted feeding | High fat diet | 4 days | NA | ad libitum vs time-restricted feeding, 4 days | Fig. 1, 2, 3<br>Ext. Data Fig. 1 and 2 | Supplementary Data 1, 2, 3 |
| Exp2 | Long-term time-restricted feeding | High fat diet | 12 weeks | NA | ad libitum vs time-restricted feeding, 12 weeks | Fig. 1 and 2<br>Ext. Data Fig. 1 and 2 | Supplementary Data 1, 2, 3 |
| Exp3 | *Dhodh* knockdown | CD/HFD | 4 weeks | NA | AAV8-ctrl vs AAV8-sh*Dhodh*, 6 weeks | Fig. 3 | NA |
| Exp4 | *Dhodh* knockdown | High fat diet | 4 weeks | NA | ad libitum vs time-restricted feeding; AAV8-ctrl vs AAV8-sh*Dhodh*, 6 weeks | Ext. Data Fig. 3 | NA |
| Exp5 | Pharmacokinetics 1 | CD | NA | single dose | IP BAY2402234 ZT0 and ZT12 | Ext. Data Fig. 4 | NA |
| Exp6 | Pharmacokinetics 2 | CD | NA | single dose | IP BAY2402234 ZT0 and ZT12 | Ext. Data Fig. 4 | NA |
| Exp7 | Early effect of BAY2402234 ZT0 | High fat diet | 4 days | daily for 4 days | VEH vs BAY2402234 at ZT0 | Fig. 4<br>Ext. Data Fig. 4 | Supplementary Data 4 |
| Exp8 | Early effect of BAY2402234 ZT12 | High fat diet | 4 days | daily for 4 days | VEH vs BAY2402234 at ZT12 | Fig. 4<br>Ext. Data Fig. 4 | Supplementary Data 4 |
| Exp9 | Time dependent effect of BAY2402234 treatment | High fat diet | 12 weeks | every other day from week 8 to 12 | VEH vs BAY2402234 vs BAY2402234+uridine at ZT0 or ZT12 | Fig. 5 and 6<br>Ext. Data Fig. 5, 6 and 7 | Supplementary Data 5 |

